# Uncoupling of the Diurnal Growth Program by Artificial Genome Relaxation in *Synechocystis* sp. PCC 6803

**DOI:** 10.1101/2021.07.26.453758

**Authors:** Anna Behle, Maximilian Dietsch, Louis Goldschmidt, Wandana Murugathas, David Brandt, Tobias Busche, Jörn Kalinowski, Oliver Ebenhöh, Ilka M. Axmann, Rainer Machné

## Abstract

In cyanobacteria DNA supercoiling varies over the diurnal light/dark cycle and is integrated with temporal programs of transcription and replication. We manipulated DNA supercoiling in *Synechocystis* sp. PCC 6803 by CRISPRi-based knock-down of gyrase subunits and overexpression of topoisomerase I (TopoI), and characterized the phenotypes. Cell division was blocked, most likely due to inhibition of genomic but not plasmid DNA replication. Cell growth continued to 4-5x of the wildtype cell volume, and metabolic flux was redirected towards glycogen in the TopoI overexpression strain. TopoI induction initially lead to down-regulation of GC-rich and up-regulation of AT-rich genes. The response quickly bifurcated and four diurnal co-expression cohorts (dawn, noon, dusk and night) all responded differently, in part with a circadian (*≈* 24 h) pattern. A GC-rich region *−* 50 bp of transcription start sites is differentially enriched in these four cohorts. We suggest a model where energy- and gyrase-gated transcription of growth genes at the dark/light transition (dawn) generates DNA supercoiling which then facilitates DNA replication and initiates the diurnal transcriptome program.

## Introduction

*In vivo*, the DNA double helix exists in a torsionally strained and underwound state, often denoted as “negative DNA supercoiling”. A homeostatic feedback system of DNA supercoiling is coupled to differential expression of large gene groups in many different bacterial species [1, 2, 3, 4, 5, 6]. The general picture (Fig. 1A) is that DNA supercoiling is high during times of high metabolic flux, such as during exponential growth, and required to express rRNA and GC-rich growth genes and allow for DNA replication [7]. The ATP/ADP ratio has direct effects on DNA supercoiling [8, 9, 10]. Supercoiling would allow an analog modulation of transcription factor and polymerase binding [11] and thereby qualifies as a potential mechanism for the long known and recently re-discovered monotonous relation of rRNA and growth gene expression with growth rate[12, 13, 14]. However, the relation of RNA transcription and DNA replication to DNA supercoiling is mutual and complex [7]. According to the twin-domain model of transcription-dependent supercoiling (Fig. 1B), negative supercoiling accumulates upstream and positive supercoiling downstream of polymerases [15], leading to cooperative and antagonistic long-range effects between transcription loci [16] (Fig. 1C). Strong transcriptional activity requires downstream activity of gyrase to set the elongation rate and avoid polymerase stalling [17, 18, 19] and upstream activity of topoisomerase I to avoid R-loop formation and genome instability [20, 21]. Such cooperative long-range effects can underpin temporal expression programs; locally in the *leu* operon [22, 23] and globally as a spatio-temporal gradient along origin-terminus axis of the *Escherichia coli* (*E.coli*) genome [24].

**Figure 1.**
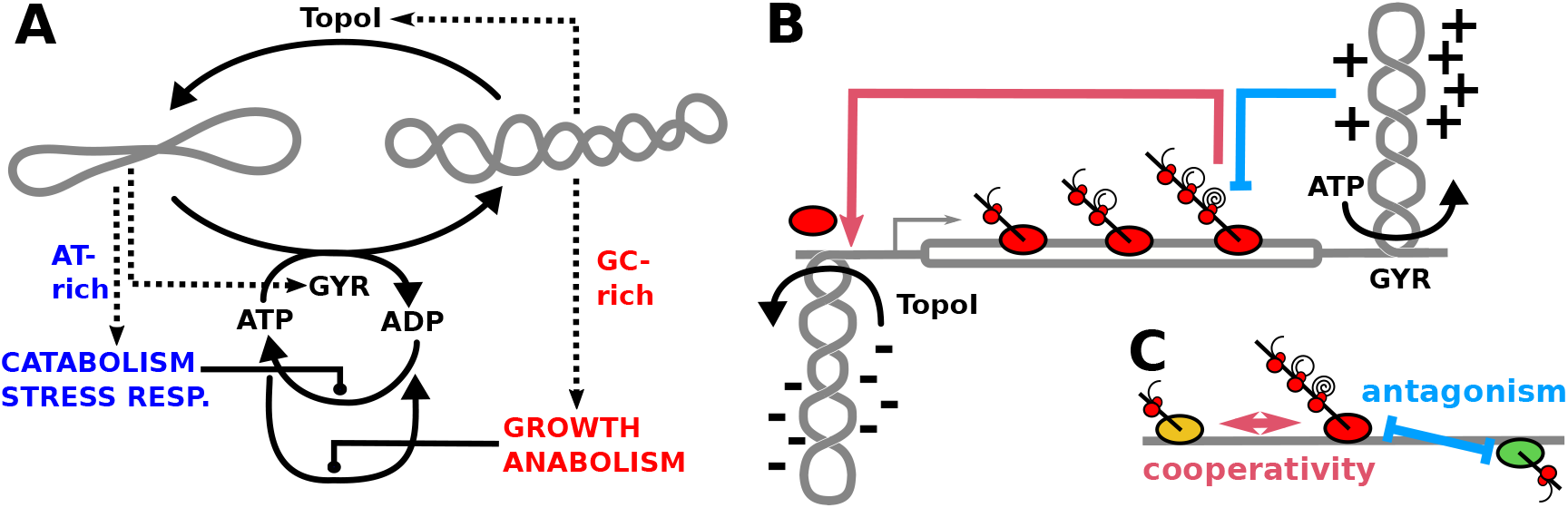
DNA Supercoiling & Transcription: Homeostasis and *Twin-Domain* Models. **A**: Global homeostasis of supercoiling by direct feedback on expression of topoisomerases (GYR: Gyrase holoenzyme; TopoI: topoisomerase I) and GC-rich anabolic/growth genes and AT-rich catabolic and stress-response genes. **B**. Transcription-dependent supercoiling downstream (positive) and upstream (negative) of an RNA or DNA polymerase, widely known as the *twin-domain* model. Gyrase-activity downstream can prevent polymerase stalling and induce transcriptional bursts while TopoI activity upstream prevents R-loop formation. **C**: The torsional stress exerted by transcription can lead to long-distance cooperative and antagonistic effects, and gene order on the chromosome can underlie temporal programs of gene expression.

In cyanobacteria, this system forms an integral part of the diurnal (light/dark cycles) changes in metabolism and transcription, and is integrated with the output of the cyanobacterial circadian clock [25, 3]. Supercoiling of chloroplast genomes, the endosymbiotic descendants of cyanobacteria, was observed to fluctuate with the diurnal light/dark (LD) cycle [26], and plants encode for gyrase enzymes [27]. Cyanobacteria themselves traverse through a well defined transcriptional program during diurnal LD cycles in several species [28, 29, 30, 31]. Mori and Johnson first suggested [32] that diurnal DNA supercoiling may be involved in the genome-wide nature of diurnal transcription in cyanobacteria. Genome compaction was found to fluctuate with LD cycles in *Synechococcus elongatus* PCC 7942 [33], and an endogenous plasmid showed diurnal fluctuations of DNA supercoiling [25]. The temporal transcriptome program during entrained circadian cycles correlated with plasmid supercoiling states and depended on gyrase activity [3]. In *Synechocystis sp.* PCC 6803 (hereafter abbreviated as *Synechocystis*), cold, heat and salt stress all lead to similar changes in the transcriptome and all were enhanced by treatment with the gyrase inhibitor novobiocin (NB) [4]. Gene groups with coherent response to stress and NB overlapped significantly with diurnal co-expression cohorts [34]. Supercoiling of the endogenous plasmid pCA2.4_M increased within 30 min after the transition to light phase, and continuously decreased during 12 h and further during a prolonged 24 h dark phase [35]^1^.

In this work, we manipulate DNA supercoiling in *Synechocystis* by inducible overexpression [36] and CRISPRi-based knock-down [37] of the key genes involved in modulation of supercoiling: topoisomerase I (gene: *topA*), and gyrase (subunit genes: *gyrA* and *gyrB*. We can confirm both, the homeostasis and transcription-dependent supercoiling models in *Synechocystis*. All manipulations that should decrease supercoiling lead to a strong pleiotropic phenotype, where cell division is blocked but cell volume growth continues. Especially *topA* over-expression induces overflow metabolism and glycogen production, and uncouples the diurnal transcription program. Metabolism and transcriptome appear to be locked in a state between late night and early day.

## Results and Discussion

### Artificial Genome Relaxation Blocks Division but not Growth

#### Manipulation of Gyrase and Topoisomerase Expression

To study DNA supercoiling in *Synechocystis*, we used the dCas9-mediated CRISPR-interference system [37] to repress (knock-down) transcription of topoisomerase I (gene *topA*, slr2058), and gyrase subunits *gyrA* (slr0417) and *gyrB* (sll2005), or *gyrA* and *gyrB* simultaneously, yielding strains named <gene>^KD^. Additionally, we constructed a tunable expression plasmid pSNDY [36] where a copy of the native *Synechocystis topA* is under the control of a rhamnose-inducible promoter, strain topA^OX^ (Tab. 1). All six strains were induced with anhydro-tetracycline (aTc) and rhamnose and cultured for five days, then harvested for quantification of cell dry weight, ATP, glycogen, and plasmid supercoiling. Reverse transcription quantitative PCR (RT-qPCR) verified the functionality of our inducible genetic constructs (Fig. S1A). All knockdown strains showed an abundance reduction to 8 %–15 % of the wild-type level, and *topA* induction was ≈30 fold. We also measured all three transcripts in all strains and observed compensatory upregulation and downregulation of the non-manipulated supercoiling enzymes, a first verification of the homeostatic control model (Fig. 1A) in *Synechocystis*.

**Table 1.**
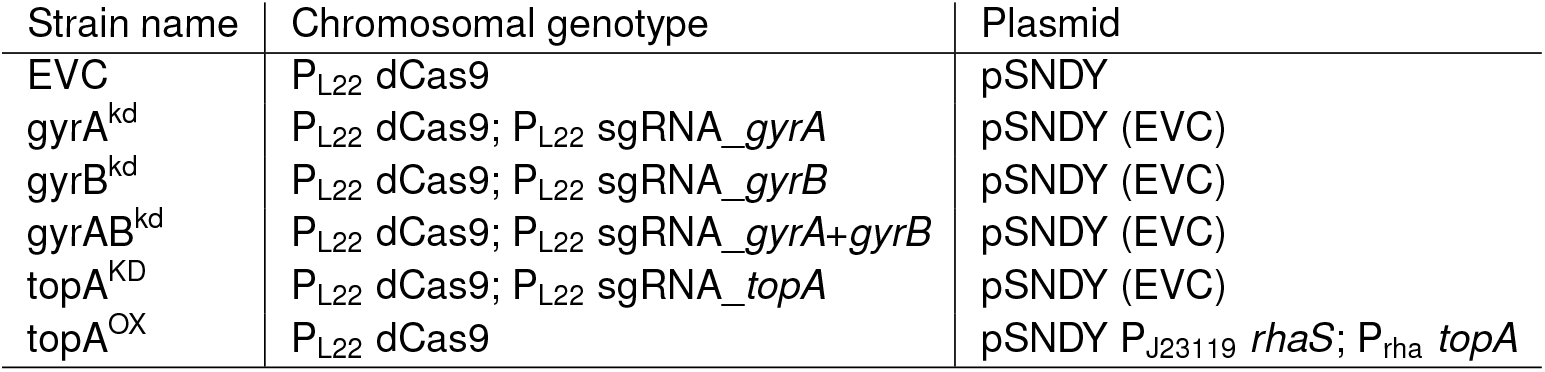
Overview of strains investigated in this work. The parental strain for all strains listed here (*Synechocystis* sp. PCC 6803 encoding aTc-inducible dCas9) was a gift from P. Hudson and L. Yao [37]

**Table 2.**
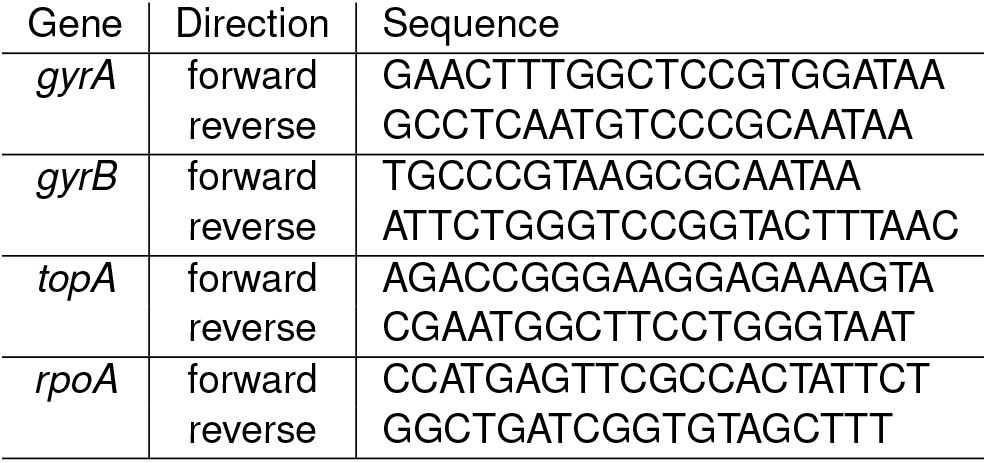
qPCR Primers.

**Table 3.**
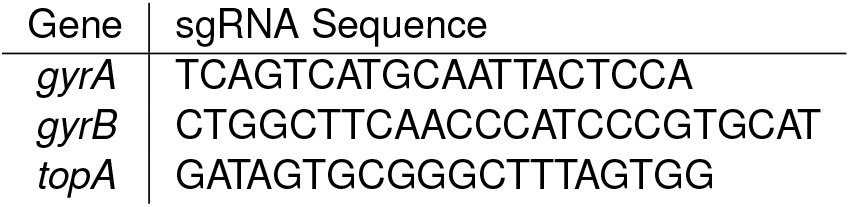
sgRNA Sequences.

#### Cell Volume Growth, Adenosine and Glycogen Content

Initially, all cultures showed comparable growth (Fig. 2A). After three days all strains except topA^KD^ grew slower than the EVC; and topA^OX^ showed the strongest growth defect. The cell dry weight (CDW) at harvest time correlated with the final OD_750_ of the cultures (Fig. 2A), but was relatively higher for the EVC and topA^KD^ strains. Cell volume distributions of the EVC and topA^KD^ strains showed a transient small increase (≈10%) on the first day of cultivation and were stable thereafter (Fig. 2B). In contrast, cell volumes of the gyr^kd^ and topA^OX^ strains increased over time, from `4 fL–5 fL to 12 fL–15 fL` after four days of cultivation. Total cell numbers increased only slightly. Thus, strains where gyrase subunits were knocked down or topoisomerase I overexpressed showed inhibition of cell division but not of cell growth. Pigmentation was strongly affected. Cultures of the topA^OX^ strain appeared pale and gyr^kd^ strains blue, compared to their uninduced state and to EVC. Absorption spectra (Fig. 2C, S1B) confirmed an overall decrease of all pigments in topA^OX^. The gyr^kd^ strains showed a stronger decrease at chlorophyll-specific wavelengths than at phycocyanin-specific wavelengths, explaining their blue appearance.

**Figure 2.**
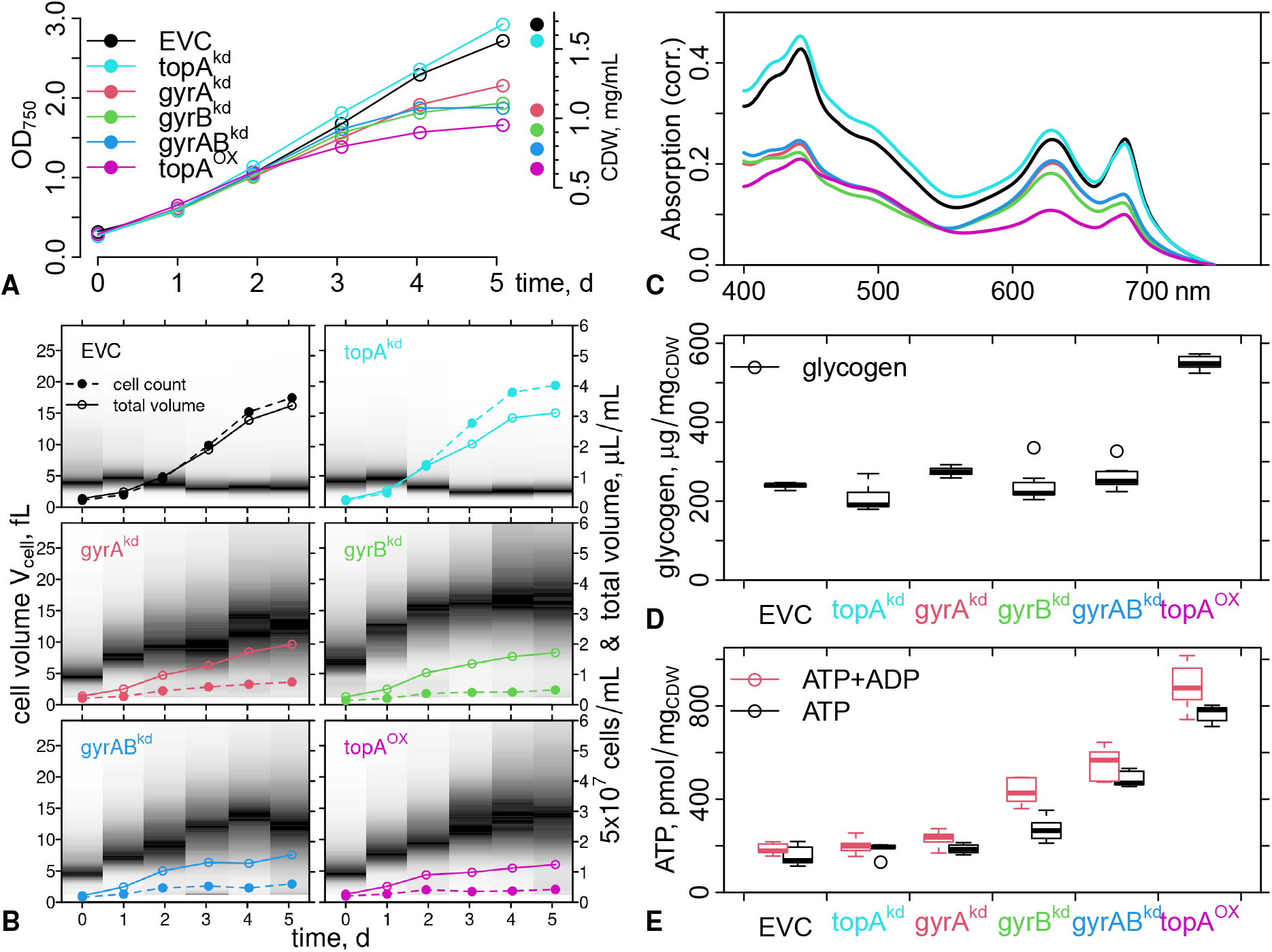
Batch Culture Endpoint Measurements. Overexpression and knock-down strains of this study where grown for 5 days in BG11 medium supplemented with all required antibiotics, and all inducers for the plasmid constructs in each experiment (100 ng/mL aTc, 1 mM L-rhamnose). **A:** The optical density at 750 nm (OD_750_) was measured daily and cell dry weight (CDW) determined directly after the last measurement on day 5. **B:** The cell volume distribution was measured daily in the CASY cell counter and plotted as a gray-scale gradient (black: more cells at this volume). **C:** Absorption spectra after the harvest on day 5. See Figure S1B for spectra at inoculation time. The absorption at 750 nm was subtracted from each spectrum. **D:** Glycogen content at harvest time was determined by a colorimetric assay after harvest, and boxplots of 18 technical replicates (3 samples, each measured 3x in 2 assays) are shown. **E:** ATP and ATP+ADP contents at harvest time were determined by a luciferase-based assay, and boxplots of six technical replicates (3 samples and 2 measurements) are shown.

All knock-down strains showed glycogen levels similar to the EVC, with 25 % of the total CDW (Fig. 2D), which is consistent with literature values for the wild type [38]. In contrast, topA^OX^ showed more than twice as much glycogen with 55 % of CDW. Values of up to 60 % are reported for nitrogen-starved cells [39, 40]. The topA^OX^ strain accumulated more than four times as much ATP+ADP as the EVC (Fig. 2E). gyrB^kd^ and gyrAB^kd^ accumulated about twice as much ATP+ADP as the EVC; topA^KD^ and gyrA^kd^ showed no difference to the EVC control. Thus, the strains repressing the ATPase subunit of gyrase, gyrB^kd^ and gyrAB^kd^, had slightly elevated levels of ATP+ADP.

#### Hypernegative Plasmid Supercoiling in the topA^OX^ Strain

Agarose gel electrophoresis in the presence of chloroquine (CQ) was used to analyze plasmid supercoiling at harvest time (5 d) of the batch cultures, and interpreted as outlined previously [35]^2^. The gels show three sets of topoisomer bands (Fig. S1D). These likely stem from the three annotated small plasmids of *Synechocystis*, pCA2.4_M, pCB2.4_M and pCC5.2_M. Electropherograms of the two smaller plasmids indicate that only strains gyrA^kd^ and gyrAB^kd^ showed plasmid relaxation (Fig. 3A). In the gyrB^kd^ and topA^OX^ strains plasmids appeared to have a higher level of DNA supercoiling. We could not extract plasmids from the topA^KD^ strain. Increased plasmid supercoiling in topA^OX^ could results from a long-term adaptation and compensatory up-regulation of gyrase subunits. We thus tested plasmid supercoiling as a time series after inoculation in fresh medium with and without the inducer (Fig. S2). The gel run time was increased to also separate topoisomers of pCC5.2_M. All three plasmids were more relaxed (less negative supercoiling) after 3 h of growth (Fig. 3B,C). Already after 8 h the trend had reversed, and at 20 h plasmids were more supercoiled than at time 0 h and in the uninduced control time series (Fig. S2E). Then plasmids became further supercoiled to an extent where topoisomers were not separable anymore. *In vivo*, hypernegative supercoiling of plasmids has been observed in topoisomerase I-deficient *E. coli* strains [41]. *In vitro*, it can be generated by gyrase and transcription [42] in agreement to the twin-domain model of transcription-dependent supercoiling (Fig. 1B). Here, the compensatory upregulation of gyrase genes apparently had a stronger effect on plasmids than the induced topoisomerase I overexpression. Plasmids and phages often harbor gyrase binding sites [43] and such sites could contribute to this phenomenon.

**Figure 3.**
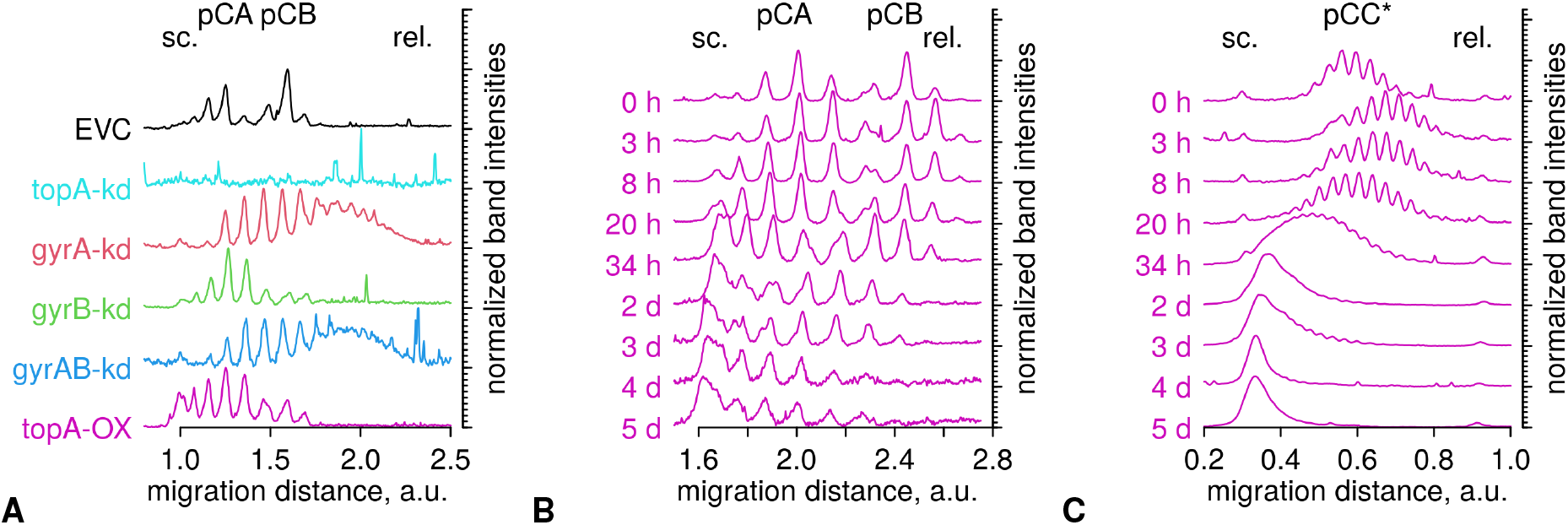
Plasmid Supercoiling. **A:** Baseline-corrected electropherograms from CQ agarose gels (Fig. S1D) of samples from the cultures shown in Figure 2, taken at harvest time (5 d). At 20 µg mL*^−1^* CQ, originally (*in vivo*) more relaxed (rel.) plasmids migrate further (higher migration distance) than more negatively supercoiled (sc.) plasmids [35]^2^. Two distinct plasmid topoisomer distributions can be distinguished in the EVC, and we assume the less far migrated bands to correspond to the longer plasmid pCA2.4_M, and the further migrated bands to the shorter plasmid pCB2.4_M. **B**: same as (A) but samples taken as a time series from the topA^OX^ strain, induced with rhamnose at 0 d (Fig. S2). **C:** as (B) but for a population of topoisomers from a larger plasmid which could be separated by running the gel longer than the gel analyzed in (A). We tentatively assign these bands to the endogenous plasmid pCC5.2_M, based on its relative migration distance.

#### Reduction of rRNA Abundances in gyr^kd^ Strains

To test effects on transcription, we selected three strains, gyrA^kd^, gyrB^kd^ and topA^OX^, and the EVC for RNA-seq analysis. Strains were grown in triplicate and harvested for flow cytometry, RNA extraction and RNA-seq analysis 5 d after induction and 3 d after a culture dilution step (Fig. S3A). Flow cytometry confirms the growth phenotype (Fig. 4A, S4): forward scatter (FSC) was increased in all strains, and most in topA^OX^. Side scatter (SSC) additionally revealed two cell populations in topA^OX^ which could reflect 8-shaped cells in division. Total nucleic acid content increased with cell size. Total RNA composition and the relative abundances of rRNA and mRNA were analyzed by capillary gel electrophoresis (Fig. S3B,C, S5). Ribosomal RNA species were strongly reduced in the gyrA^kd^ and gyrB^kd^ strains and less reduced in topA^OX^ (Fig. 4B), except for the 600 bp fragment of the 23S rRNA (Fig. S6C). In *E. coli*, transcription of ribosomal RNA depends on downstream gyrase activity [44], mediated by strong gyrase binding sites [19]. Such an interaction would be directly (locally) affected in the gyr^kd^ strains. In contrast to plasmids, the upregulation of gyrase in the topA^OX^ strain could apparently not locally compensate for global genome relaxation by topoisomerase I, thus also reducing the relative rRNA content in this strain.

**Figure 4.**
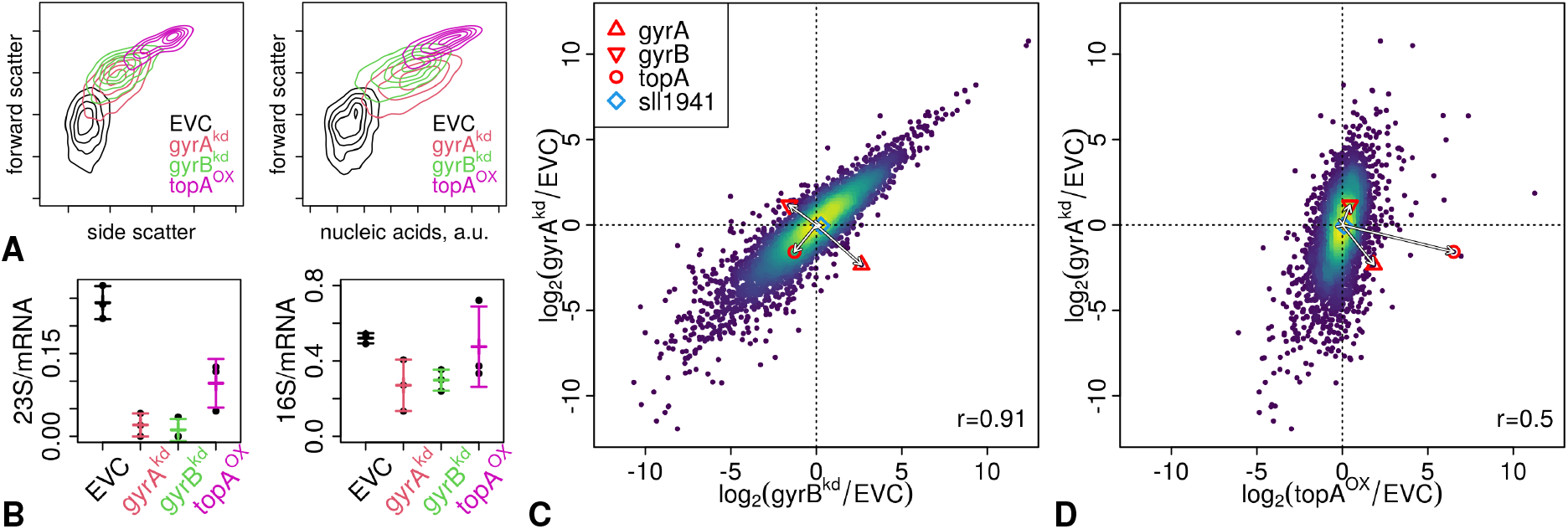
rRNA and mRNA Abundances in Induced Strains. **A:** Flow Cytometry confirms the cell growth phenotype. The natural logarithms of forward scatter, side scatter and nucleic acid stain Syto9 were calculated and 2D distributions plotted as contour plots; see Figure S4 for all data and details. **B:** Relative abundances of the unfragmented 23S (B) and the 16S (C) rRNA species were calculated from the electropherograms of capillary gel electrophoresis of extracted RNA (Fig. S5). Figure S6 shows all rRNA species. Mean and standard deviation of the three replicates are indicated as error bars. **C & D:** Relative expression changes of coding genes were derived as the log_2_ ratio of RPKM normalized read-counts in the induced over-expression or knock-down strains to values in the control strain (EVC) and compared between the three different strains by 2D histograms (yellow: highest and purple: lowest local density of genes). The Pearson correlations (r) are indicated in the bottom right corner. **C**: gyrA^kd^ (y-axis) *vs.* gyrB^kd^ (x-axis) strains. **D**: gyrA^kd^ (y-axis) *vs.* topA^OX^ (x-axis). The induction/repression and the homeostatic responses of *gyrA*, *gyrB* and *topA* are highlighted by arrows from the origin to indicate the direction of change. The gene *sll1941* is a homolog or paralog of *gyrA* (blue diamond) and its transcript showed no response in either experiment.

#### Consistent Changes in mRNA Abundances & Homeostatic Regulation of Supercoiling Enzymes

The same RNA samples were further processed (rRNA species depleted) and sequenced on the Illumina platform, and transcript abundances relative to the EVC evaluated with DESeq2 [45]. All strains showed overall similar expression changes, but the extent was lower in topA^OX^ (Fig. 4C,D). However, this difference could also just reflect normalization effects by the decreased rRNA content in the gyrase knock-downs. In all strains the targeted manipulation was still observable at harvest times (arrows in Fig. 4C,D), *i.e.*, *gyrA* transcripts were reduced in gyrA^kd^, *gyrB* transcripts in gyrB^kd^ and *topA* transcripts were increased in topA^OX^. The non-manipulated genes showed the compensatory response expected from homeostatic regulation, *i.e.*, *topA* was repressed in both gyr^kd^ strains, and all non-manipulated gyrase subunits were induced in all experiments. In contrast, the *sll1941* gene, annotated either as a second gyrase A subunit or as the topoisomerase IV ParC subunit, showed no response in either experiment. Thus, RNA-seq confirms the RT-qPCR results, further corroborating the homeostatic regulation model (Fig. 1A), and emphasizes the genome-wide nature of DNA supercoiling effects.

### Overexpression of *topA* Uncouples Diurnal Co-expression Cohorts

#### Transient Increase in Cell Volume and Density

To study the dynamic response to transient *topA* induction, the topA^OX^ strain was grown in a Lambda Minifor bioreactor (Fig. S7) with continuous (online) monitoring of turbidity (OD*_λ_*, Fig. S8A,B). Continuous culture dilution was initiated at OD*_λ_* ≈ 2.9 and with dilution rate *φ* ≈ 0.24 d*^−1^*. The culture stabilized around OD*_λ_* ≈ 2.7. Notably a subtle ≈ 24 h pattern of OD*_λ_* was observed in both, batch and pre-induction continuous growth phases. Then rhamnose was injected to 2 mM to induce overexpression of *topA*. The *topA* transcript was upregulated to ≈45-fold over the pre-induction level within 4 h, as measured by RT-qPCR and confirmed by RNA-seq (Fig. S11) and decreased slowly over the course of the experiment. The OD*_λ_* initially increased for 1 d post-induction, then slowly decreased. Cell dry weight (CDW) measurements were noisy but matched the OD*_λ_* signal over the sampled period (Fig. 5A, S8C). In contrast, cell numbers started to decrease immediately, and cell volumes increased (Fig. 5B). We calculated growth rates of OD*_λ_*, cell numbers and the total cell volume (Fig. 5C, S9). Cell division was not completely blocked but severely reduced to a division time of ≈10 d (*µ*_count_ ≈ 0.07 d*^−1^*). Total cell volume growth was much less affected and remained stable (*µ*_volume_ ≈ 0.18 d*^−1^*) throughout continuous culture operation until 12 d post-induction. Thus, artificial *topA* overexpression blocked cell division but not cell volume growth. OD*_λ_* growth remained highest (*µ*_OD_ ≈ 0.23 d^−1^) and stable over the first 5 d–6 d. In parallel, glycogen content increased to about 35 %–40 % of the CDW (Fig. 5A). We further noticed that sampled cells started to sediment much faster, indicating increased intracellular density. By calibrating the OD*_λ_* signal to the CDW measurements (Fig. S8C) and dividing by the total cell volume we can estimate a CDW density and this value also increased over time from 0.3 to 0.5 g_DCW_/mL_cell_ (Fig. 5D). This range is consistent with data from *E. coli* [46, 47]. However, the CDW per OD_750_ was relatively lower for the enlarged strains in the endpoint measurement (Fig. 2A), and thus, the calibration to OD*_λ_* may overestimate true CDW density. The enlarged cells also became increasingly fragile: in the CASY cell counter data a small population of varying intensity appeared at *<* 2 fL. This peak was highest at 7 d (outlier x in Fig. 5B), where cells were lysed during centrifugation in a washing step. The washing step was skipped thereafter, and the peak of small cells (dead or fragmented) remained small but increased towards the end of the continuous culture. Maximal cell volumes >20 fL were reached 10 d–15 d post-induction. From day 14 a population of smaller cells, ≈7.5 fL, appeared. On 16 d this population was the majority, and cell volume further decreased to 5 fL. Cell pigmentation recovered and the culture appeared greener again. We then switched off dilution, and the culture resumed growth, although at lower growth rates than pre-induction.

**Figure 5.**
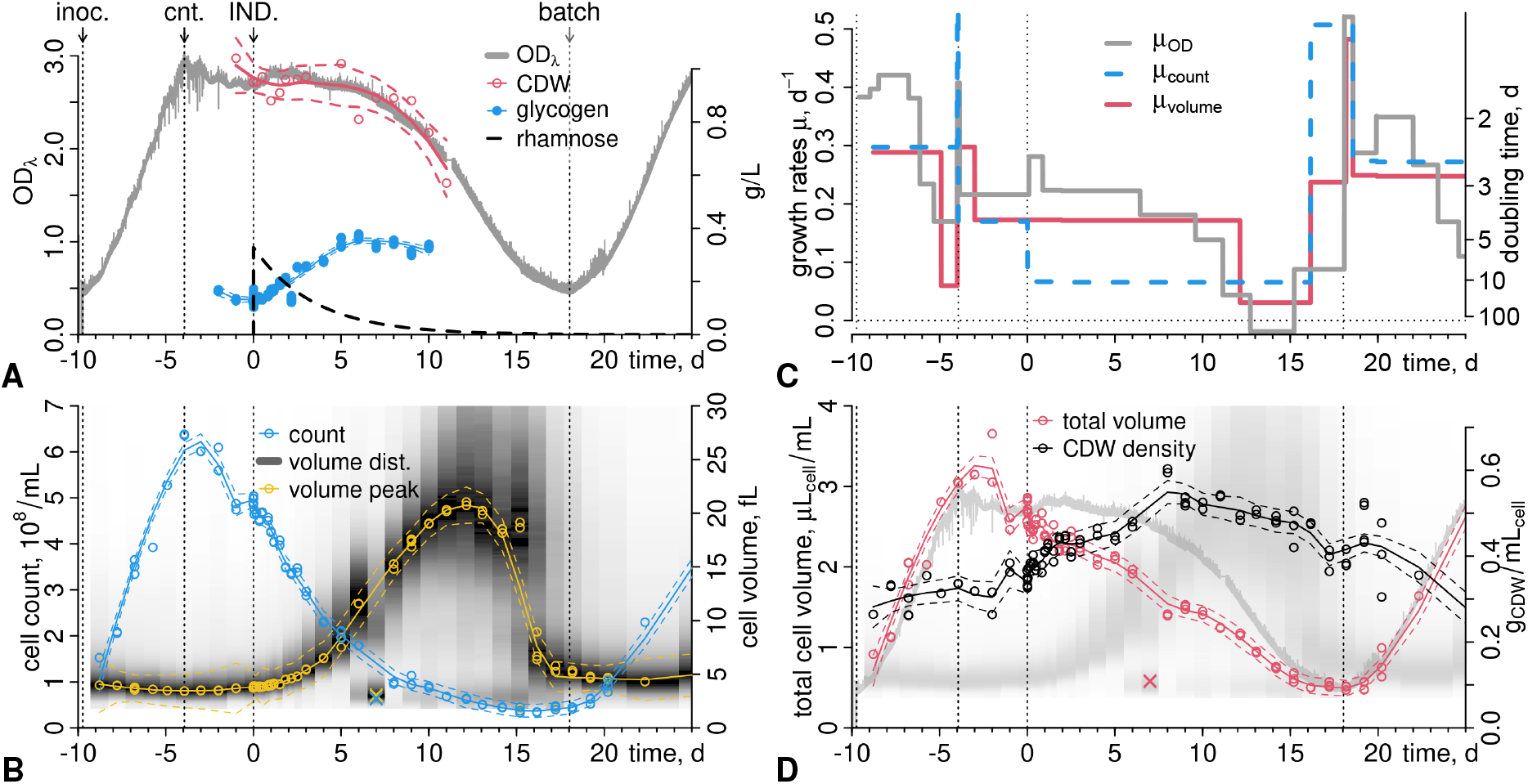
Pulsed Induction in Continuous Culture. **A:** Photobioreactor growth of the topA^OX^ strain (1 L BG11 medium, 0.5% CO_2_, illumination 90 µmol m*^−^*^2^ s^−1^ per OD_750_). Optical density was recorded online (OD*_λ_*) and post-calibrated to offline OD_750_. The arrows indicate inoc.: inoculation; cnt.: onset of continuous culture (rate *φ* = 0.01 h^−1^); `IND`.: induction of *topA* by pulse-addition of rhamnose to 2 mM (0.33 g L^−1^) at time 0 d; and batch: switch-off of dilution. The dashed black line is the theoretical wash-out curve of rhamnose (g L^−1^). Cell dry weight (CDW, g L^−1^, red) and glycogen content (g L^−1^, blue) of the culture were measured at the indicated times (points), and LOESS regressions are shown (solid lines) with 95% confidence intervals (dashed lines). **B:** Cell numbers (blue points) and volume distributions (gray scale) were recorded daily, and at higher resolution after induction, with the CASY cell counter. The peaks of the cell volume distributions are shown as yellow points. LOESS regressions with 95% confidence intervals are shown as lines. One outlier of the CASY measurement (x) was due to cell lysis during a washing step and was not included for regression. **C:** growth rates *µ* were calculated by local (piecewise) linear regressions of the OD*_λ_* (A), and cell count (B) and total cell volume (D) measurements and subtraction of the culture dilution rate (Fig. S9). **D:** The total cell volume (*V*_total_) was calculated by integrating the single cell volume distributions in (B), and the CDW density was calculated by dividing the OD*_λ_* signal, calibrated to the CDW measurements (A, Fig. S8C), by *V*_total_.

#### Upregulation of Plasmid & Growth Genes, Downregulation of Photosynthesis Genes

Samples for total RNA analysis and RNA-seq were taken 1 d and 0.5 h before induction, and then over the next 25 d in decreasing temporal resolution to roughly capture three time-scales of the response. Coding gene transcript abundances were calculated, the resulting time series clustered (Fig. 6A, S12), clusters sorted along significant overlaps with diurnal cohorts (Fig. 6C), and functional annotation enrichments (CyanoBase “categories”: Fig. 6B, S14A; gene ontology: Fig. S15) calculated. Cluster 2 (red) comprises the majority of the ribosomal protein category, enzymes of RNA and DNA synthesis, amino acid biosynthesis and the ATP synthase. Transcript abundances increased slightly over the three days after induction, with a notable circadian (≈ 24 h) pattern. Cluster 3 (yellow) comprises the majority of photosynthesis-related genes, including the two photosystems. Its transcript abundances decreased continuously. The remaining clusters showed less clear functional profiles. Clusters 1 (green) and 4 (blue) were enriched in genes with unknown function, and cluster 4 strongly enriched with transposase genes. Gene Ontology analysis additionally shows enrichments with 7 of 8 genes annotated with “DNA polymerase activity” and 60 “DNA binding”proteins (Fig. S15). Both clusters 1 and 4 showed decreased transcript abundances on the first day post-induction, then only cluster 4 increased steeply. Cluster 1 transcripts showed a subtle circadian pattern, peaking anti-phase to cluster 2 transcripts. Cluster 6 (cyan) showed a similar profile to cluster 4 and comprises transcripts that showed the strongest abundance increase. Both clusters 4 and 6, were strongly enriched for plasmid-encoded genes (Fig. S16), and both also contained significant fractions of the ribosomal protein category. Cluster 5 (gray) showed decreased abundances but had the least change over time, and was slightly enriched with “hypothetical”, “Hydrogenase” and “plasma membrane” gene annotations.

**Figure 6.**
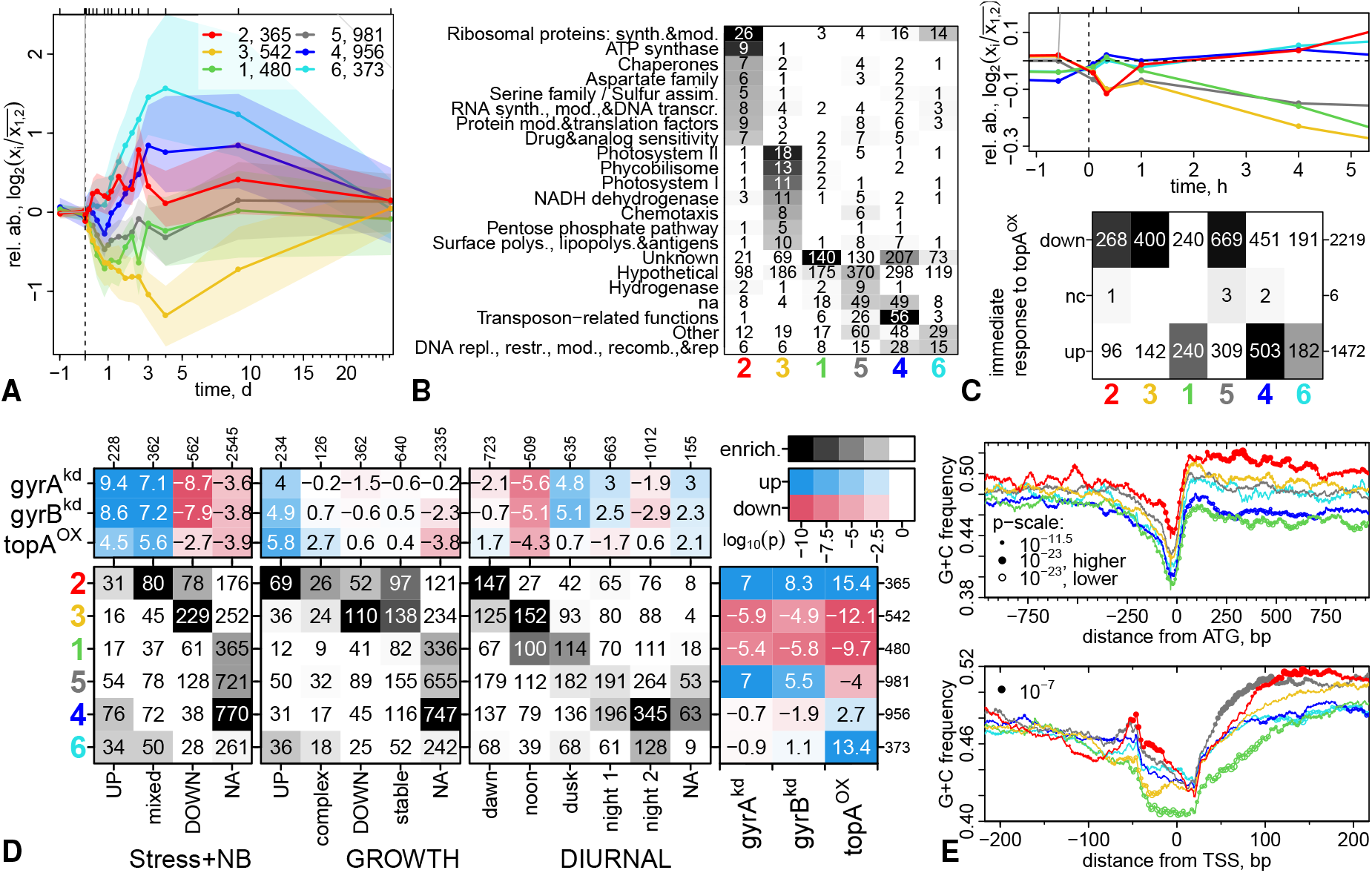
Co-Expressed Gene Cohorts. **A:** Cluster medians (colored solid lines) and 25%/75% quantiles (transparent ranges) of relative transcript abundances (rel. ab.: log_2_ ratio (TPM) to the mean TPM of two pre-induction samples); points indicate sampling times (ticks on upper axis). Cluster labels and sizes are indicated in the legend. **B:** Sorted enrichment profile of co-expression cohorts with the CyanoBase “categories” gene annotation; black field: *p* ≤ 10*^−^*^10^; white text: *p* ≤ 10*^−^*^5^, row sorting and filter: *p*_sort_ ≤ 0.01. Figure S14A shows unfiltered results for all categories. Numbers are counts of genes in both categories. **C, top:** Cluster medians as in (A) but zoomed in on the first five hours after induction and without – the quantile ranges. **C, bottom:** Cluster enrichment profile (black: *p* ≤ 10*^−^*^10^; white text: *p* ≤ 10*^−^*^5^) with genes upregulated (up), downregulated (down) or without change (nc) immediately (5 min–20 min) after induction. **D:**. Enrichment profiles (gray scale) with other published gene classifications (see text, Fig. S14B, S17)) and t-value profiles (red-blue scale) of clusters in the end-point transcriptome experiments. Blue: *t >* 0, red: *t <* 0; *p*_min_ = 10*^−^*^10^, white text: *p* ≤ 10*^−^*^5^. The text indicates total counts as in (B), or *t*-values in t-test profiles. **E:** GC-content profiles around start codons (ATG, top panel) and transcription sites (TSS, bottom panel) from S23). GC-content was calculated in 66 bp windows at each position. Point sizes scale with log_2_(*p*) from local motif enrichment tests (filled points) and deprivation (open circles) tests, and the minimal p-values are indicated in the legends. These significance points are only shown every 10*^th^* (top) or 3*^rd^* position (bottom).

#### Uncoupling the Diurnal Program at Dawn

Next, we compared the time series clusters from topA^OX^ and the endpoint transcript abundances from the topA^OX^ and gyr^kd^ strains with previously characterized co-expression cohorts (Fig. 6D). Prakash *et al.* clustered genes into three groups by their response to salt, heat and cold stress, each with or without the gyrase inhibitor novobiocin (NB) [4]. Zavrel *et al.* measured protein abundances at different growth rates in constant light conditions and presented 7 clusters [14] which we summarized into cohorts upregulated or downregulated with growth rate, with a complex response and stable proteins without growth rate-dependence (Fig. S14B). Saha et al. and Lehmann *et al.* analyzed transcriptome time series over light-dark cycles [48, 34]. Saha *et al.* did not cluster their data, thus we clustered their data into 5 cohorts and assigned them names that correspond to their order of expression during the diurnal light-dark cycle (Fig. S17). Clusters 2 (red) and 3 (yellow) show the clearest functional profiles, and are found co-expressed with opposite patterns in all tested experiments, and at dawn and noon, respectively, in the diurnal data sets. Cluster 2 was most enriched with transcripts that showed a mixed response to the applied stress conditions (`Stress+NB`) and with genes whose protein abundances correlated positively with growth rate (`GROWTH`). Cluster 3 was most enriched with transcripts downregulated in stress and whose protein abundances correlated negatively with growth rate. In the diurnal data set cluster 2 transcript abundances peaked in early day (dawn) and were followed by a peak of cluster 3 transcripts at noon. These clusters likely reflect biological co-expression cohorts and from hereon we denote cluster 2 as cohort `RB/dawn` (ribosomal proteins and biosynthesis) and cluster 3 as `PS/noon` (photosynthesis), according to their annotation and experiment enrichment profiles. Cluster 1 is enriched with proteins of unknown function. In both, stress and growth experiments, we found only enrichments in the NA classes, ie. the original papers did not report values. However, in the diurnal data set we found enrichment with genes clusters that peak at noon or after noon. We denote this cohort as UK/dusk (unknown function, expressed at dusk). The remaining clusters 5, 4 and 6 all show weak functional enrichments. Only cluster 4 is enriched with transposase genes, DNA polymerase and other DNA binding proteins. All of them, and strongest in cluster 4, show weak enrichment with transcript cohorts that peak during the dark phase. Cluster 6 is enriched in the late night cohort in the diurnal data set, and further shows some overlaps in enrichment with the `RB/dawn` cohort, thus closing the (diurnal) cycle of expression. We tentatively label cluster 4 as the `DNA/night` cohort and cluster 6 as plasmids.

To compare the time series with the endpoint transcriptome data, we calculated t-test profiles of all clusters (red/blue squares in Fig. 6D and S13). `RB/dawn`, PS/noon and `UK/dusk` showed consistent behavior in all strains, although at lower t-values for the gyrase knock-downs. `DNA/night` and `plasmids`, both enriched with plasmid genes, were upregulated only in topA^OX^ but not in the gyr^kd^ strains. Transcripts of the large cluster 5, which showed the weakest response in the time series, were upregulated only in the gyr^kd^ strains. Likewise, the diurnal co-expression cohorts from ref. [48] (Fig. S17) all show subtle but notable differences between the strains, only PS/noon was consistently downregulated.

#### Differential Regulation of GC-rich and AT-rich Genes

Genes with supercoiling-activated transcription were found to be GC-rich in both upstream non-coding and coding regions of several species, and *vice versa*, supercoiling-repressed genes are AT-rich [1, 3, 49, 6]. Thus, we aligned nucleotide sequences of all genes from the main chromosome at their start codons (ATG), calculated average nucleotide content in moving windows for each cluster, and performed a statistical enrichment test (cumulative hypergeometric distribution) at each position (Fig. 6E). The up-regulated `RB/dawn` cohort and the down-regulated `PS/noon` cohort are GC-rich downstream and upstream of the start codon. The `UK/dusk` (downregulated) and `DNA/night` cohorts (upregulated) are AT-rich. Zooming in to the immediate response after induction shows that the response of the co-expression cohorts switches within the first h post-induction (Fig. 6C). The immediate response is consistent with data from other species. Transcript abundances of both GC-rich cohorts, RB/dawn and `PS/noon`, decreased, while those of the AT-rich `UK/dusk` and `DNA/night` cohorts increased. Already 1 h post-induction, the two GC-rich and two AT-rich cohorts have bifurcated. Transcript abundances of `RB/dawn` increased while those of `PS/noon` continue to decrease. Similarly, the AT-rich UK/dusk transcripts now decreased while DNA/night transcripts continued to increase in abundance.

#### Promoter Structure: A GC-rich Discriminator?

To analyze actual promoter structures we mapped the coding genes on previously described transcription units (TU) [50], calculated average temporal abundance profiles of TUs, and clustered TU profiles by `k-means` using the average profiles of the gene-based clustering as cluster centers (Fig. S23). TU were then aligned at their transcription start sites (TSS) and again average nucleotide content profiles of clustered TU calculated (Fig. 6E). Downstream of the TSS the general differences in GC/AT content were similar to those of start codon-aligned coding genes, albeit with lower statistical power. TSS alignment reveals a distinct GC-rich peak at ca. 50 bp. This peak corresponded better to the mid-term trend of transcript abundances. Cohorts that were upregulated within the first 3 d post-induction (`DNA/night` and `RB/dawn`) had a higher, and downregulated cohorts (PS and UK/dusk) had the lowest GC-content. In summary, the immediate effect of genome relaxation was quickly overruled by other regulatory mechanisms. A candidate is the ppGpp-mediated repression of day-time transcription [51]. In *E. coli*, a GC-rich discriminator at −10 bp mediates the supercoiling-dependence of the ppGpp-mediated “stringent response” of promoters [52, 53]. Conceivably, the 50 bp site could have a comparable role in integrating the response to different regulatory inputs.

#### Genes of Interest

In Figures S18–S22, we analyze transcript abundances of a few specific gene sets: circadian clock genes (*kai*), response regulators, sigma factors, and photosynthesis and metabolic genes. However, these patterns are hard to interpret, due to the unknown function of upregulated genes and differential regulation of paralogs and enzyme complex subunits. In short: most *kai* genes were down-regulated, while *kaiC3* is upregulated with RB/dawn, followed by *kaiB2* and *kaiC2* from 3 d (Fig. 7A, S18A). KaiC3 is required for chemoheterotrophic growth in constant darkness [54]. The response regulator *rpaB* is downregulated, and *rpaA* slightly upregulated (Fig. S18B). Of the tested regulators, only *pmgA* is strongly upregulated over the first three days, with the diurnal pattern of the RB/dawn cohort. Sigma factors of unknown function [55], *sigH → segI*, were upregulated until 3 d; the stress factor *sigB* is up-regulated only at 10 d (Fig. S18C), where cells are enlarged. Photosystem, phycobilisome, and carboxysome genes are predominantly downregulated (Fig. S19–S20). In contrast, RuBisCo and genes of the carbon concentrating mechanisms are predominantly upregulated over the first three days (Fig. S21). The glycogen degrading enzyme *glgP1* was strongly downregulated; glucose-1-phosphate adenylyltransferase *glgC* was upregulated, and glycogen debranching enzymes *glgX*/*X2* peaked at 3 d (Fig. S22A). NAD synthesis genes are generally upregulated, and we find strong differential response of certain subunits of the NADH dehydrogenase (Fig. S22B,C). Together with the upregulation of the redox and photomixotrophic growth regulator *pgmA* [56], these responses point to changes in redox metabolism. The thioredoxin *trxM1* is strongly down-regulated, and *trxA* has only a short and low peak at 2 d–3 d (Fig. S22D).

**Figure 7.**
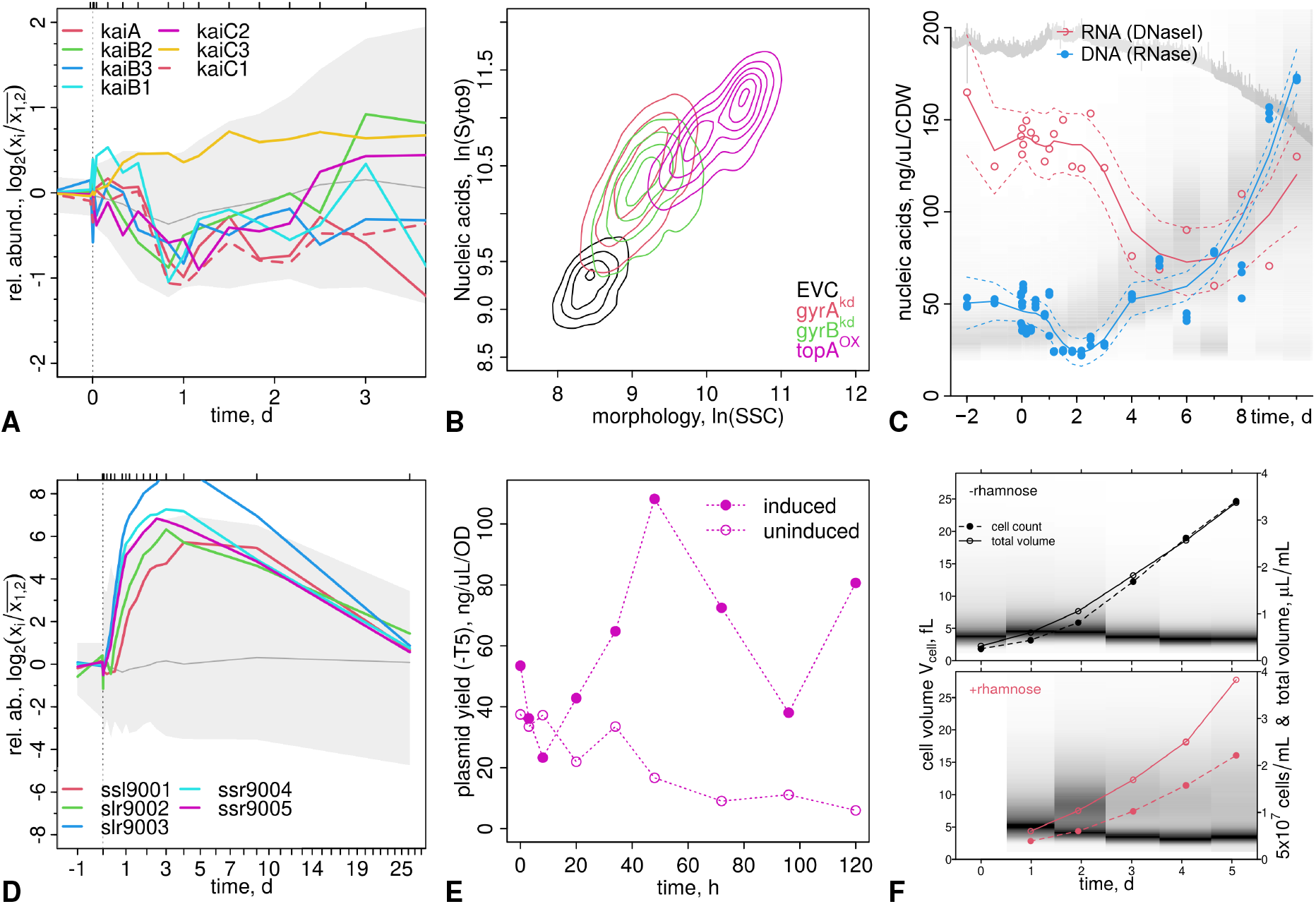
Explaining the Phenotype? **A**: Relative transcript abundances of the Kai circadian clock genes in the topA^OX^ time series, only *kaiC3* is upregulated over the first three days post-induction. The gray background shows the 92.5 % and 7.5 % quantiles of all data as a reference. **B:** Flow cytometry of batch culture endpoint samples, as Fig. 4A but for nucleic acid content (Syto9 marker) *vs.* the side scatter signal, an indicator of cell morphological features. **C:** RNA and DNA extraction yields in the topA^OX^ time series experiment, divided by the OD*_λ_* signal calibrated to CDW. OD*_λ_* and cell volume distributions (Fig. 5A,B) are shown (gray) as reference. Lines show a LOESS regression with 95% confidence interval. **D:** Relative transcript abundances (as in (A)) of genes from the plasmid pCC5.2_M. **E:** Plasmid extraction yields in the plasmid supercoiling time series (Fig. 3) from the induced and uninduced cultures. **F:** Cell volume and count development after re-induction of of cells isolated after harvest from the topA^OX^ time series experiment in fresh BG11 medium.

#### Reversible Inhibition of DNA Replication Explains the Growth Phenotype

Manipulation of DNA super-coiling has highly pleiotropic effects, and can also interfere with DNA replication and lead to DNA damage [57]. Here, we observed a block of division while cell growth continued. Could an inhibition of genome replication be involved? In endpoint RNA-seq experiments relative rRNA abundances were decreased but the total nucleic acid content (Syto-9 stain) increased with cell size and morphology (Fig. 4A, 7B, S4). To differentiate nucleic acids, we extracted RNA and DNA from the topA^OX^ time series experiment (Fig. 7C). Total DNA extraction yields (per CDW) dropped to about 50 % in the three days post-induction. From 4 d post-induction, DNA yields increased steeply; but increased plasmid-derived transcript abundances (Fig. 7D, S16) and increased plasmid extraction yields (Fig. 7E, S2B,C) suggest that this was due to increased plasmid replication. Thus, the most parsimonious explanation for the division block phenotype is impaired DNA replication of the genome, while the copy numbers of small plasmids increased during the late stages of cell volume growth. A block of genome replication could be due to permanent DNA damage. The volume growth and division block phenotype was reversible. All strains recovered readily when re-inoculated in fresh medium without the inducer (Fig. S1C). In the continuous culture experiment, cells recovered as the inducer was washed out (Fig. 5A). Re-inoculation with and without inducer showed that in some cells our constructs remained intact and induced cells again grew in volume, while other cells were refractive to induction, and overgrew the division-blocked cells (Fig. 7F, S10). Thus, we conclude that the block in genome DNA replication was not due to irreversible or lethal genome damage.

## Conclusion

We presented a comprehensive characterization of DNA supercoiling effects in *Synechocystis*. Several models of the role of this regulatory system in bacterial cell biology can be confirmed in this species. The enzymes directly responsible for supercoiling are under homeostatic control [60] (Fig. 1A). Increased gyrase levels and transcription conspire to hyper-supercoil plasmids in the topA^OX^ strain, in agreement with the twin-domain model of transcription-dependent supercoiling [15, 42] and suggesting the presence of gyrase binding sites 1. [43] in the small endogenous plasmids of *Synechocystis*. Differential transcription of GC-rich and AT-rich genes [1, 3, 49, 6] was observed immediately (5 min–20 min) after induction of *topA* over-expression. Already after 60 min the pattern had changed. The compensatory upregulation of gyrase and other regulatory mechanisms, such as the ppGpp mediated repression of day time transcription [51], may underlie this bifurcation of the response; potentially mediated by a GC-rich “discriminator” 50 bp upstream of TSS. Large gene cohorts developed similarly over the first three days post-induction, and these revealed a tight link to the diurnal transcription program, consistent with results from *Synechococcus elongatus* PCC 7942 [3].

Cooperative and antagonistic long-range effects of transcription *via* local accumulation and diffusion of DNA torsional strain [16] (Fig. 1B,C) underpin temporal expression sequences in operons [22, 23], and genome-wide along the origin-terminus axis during *E. coli* growth phases [24]. In analogy to these models, we suggest a simple hypothesis of supercoiling-mediated cooperativity during the diurnal transcription program in cyanobacteria (Fig. 8). Overall, genomic supercoiling is decreased in our strains, despite availability of light-derived energy. This leads to upregulation of the night-expressed gene cohort (`DNA/night`), incl. the *kaiC3* gene [54] despite the constant light conditions. The availability of light-derived energy drives transcription of ribosomal RNA and growth genes around dawn (RB/dawn) from relaxed DNA, *e.g. via* low levels of ppGpp and high GTP. But this process is gated or modulated by local activity of gyrase, bound to putative gyrase binding sites downstream of the affected transcription units. In wildtype conditions, this transcription-dependent supercoiling would increase genome-wide supercoiling and thereby directly facilitate both DNA replication between dawn and noon [61] and advancement of the diurnal transcription program to the GC-rich PS/noon gene cohort. In our strains, genome-wide supercoiling can not accumulate and these latter steps are blocked. Cells are stuck in a state between late night and dawn.

**Figure 8.**
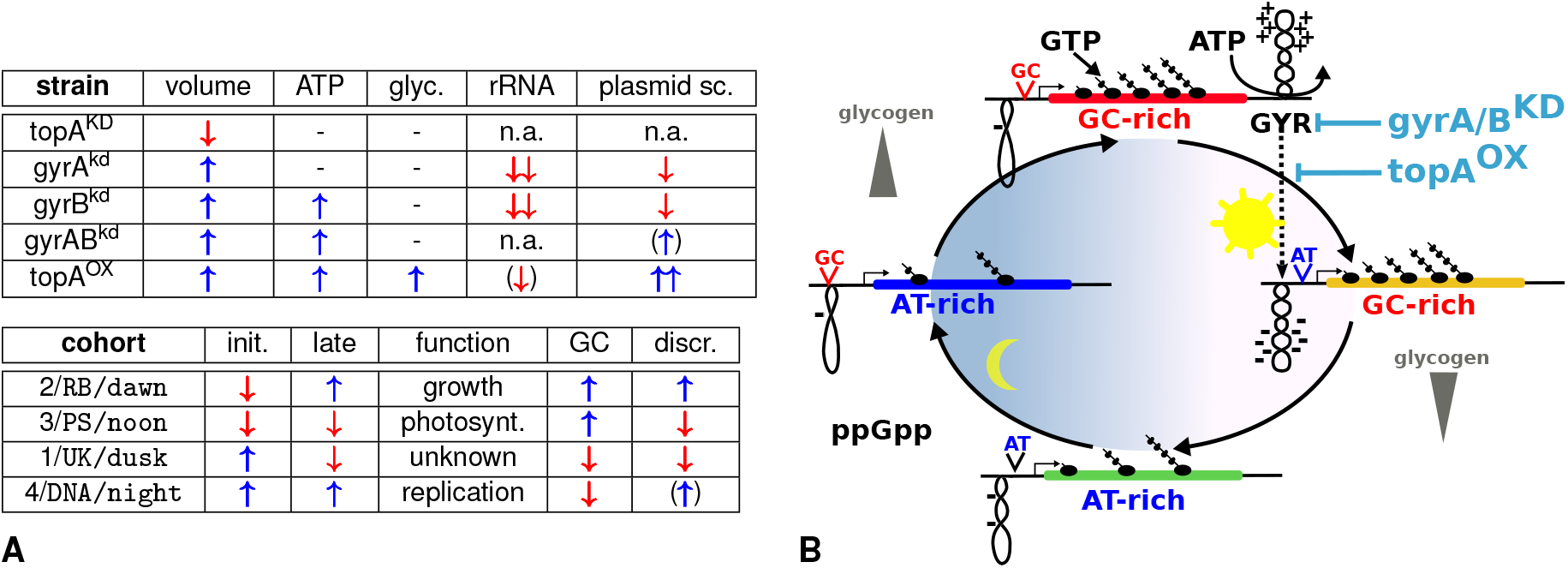
Summary and Model. **A:** Summary tables of physiological characteristics of the tested strains (top) and the co-expression cohorts observed after topA^OX^ induction (bottom). The top table indicates changes in cell volume, ATP+ADP, glycogen, rRNA content and plasmid supercoiling. The bottom table summarizes enriched properties of the co-expression cohorts (clusters): the initial and late response of transcript abundances, functional enrichments, overall GC content and the presence of a putative GC-rich discriminator at 50 bp upstream of the TSS. **B:** Development of an Integrative Model: at dawn energy from photosynthesis becomes available, allowing for transcription of GC-rich growth genes (RB/dawn), gyrase activity downstream of these genes can gate their transcription, *e.g.* tune bursting frequency [58] or set the elongation rate [44]. This is inhibited in the gyrase knock-downs and not inhibited or even enhanced through gyrase upregulation in the *topA* over-expression experiments. The combination of transcription and gyrase activity leads to genome-wide increase in negative supercoiling. The accumulated torsional strain can then be channeled into transcription of the second GC-rich cohort, `PS/noon`. This latter process is inhibited in all strains. In normal conditions topoisomerase I could then be involved in resolving R-loops [20] that may have been generated by the dawn/noon program. During night-metabolism ppGpp signaling [51] activates the night transcription program, at low levels of transcription [31], and supercoiling globally decreases. Clock-mediated glycogen mobilization at the end of the night [59] could kick-start the RB/dawn program already before onset of light [48].

To test this model, our strains should next be studied in this natural diurnal context, that is, in cultures grown in regular day/light cycles, and interference (induction) initiated at different times of the cycle. Further development of artificial regulation of supercoiling enzymes, towards a full de-construction and re-construction of the homeostatic feedback system in one strain, would provide a highly versatile tool to untangle the pleiotropic effects of DNA supercoiling on transcription, replication and cell growth. And finally, topoisomerase I overexpression caused an increase of cellular glycogen content of up to 60 % of the cell dry weight. This phenotype could be exploited for synthesis of high value products or directly as a fermentation substrate for yeast [40]. Thus, further development of such strains may be of biotechnological value.

## Data Availability

The clustering and time series data from the topA^OX^ strain (both as raw abundances in TPM and as the log2 ratios to the mean of two pre-induction values, as plotted in this manuscript), and endpoint measurements (log2 ratio of abundances in the gyrA^kd^, gyrB^kd^ and topA^OX^ strains to the EVC strain) are available as Datatable_S1.tsv. *** The raw sequencing reads will be made available (*e.g.* SRA) upon publication. ***

## Funding

RM was funded by the *Deutsche Forschungsgemeinschaft* (DFG), grants AX 84/4-1 & STA 850/30-1 (COILseq) and EXC-2048/1–project ID 390686111 (CEPLAS). MD, DB and TB were financially supported by the CLIB Competence Centre Biotechnology (CKB) funded by the European Regional Development Fund ERDF (EFRE-0300096 and EFRE-0300095).

## Acknowledgments

The parental strain *Synechocystis* sp. PCC 6803, encoding aTc-inducible dCas9, and the gyrB^kd^ strain were a gift from P. Hudson and L. Yao. We are grateful to Nic Schmelling for critical discussion of the manuscript.

## Materials and Methods

### Strains and Plasmids

The *Synechocystis* parent strain used for all genetic alterations was kindly provided by Lun Yao and Paul Hudson, and based on a strain they had obtained from Dr. Martin Fulda (Göttingen, Germany). It contains a TetR cassette, as well as dCas9 under the promoter PL_22_, inducible with anhydrotetracycline (aTc), at the genomic insertion site psbA1. Construction of this strain was described in ref. [37]. For overexpression of genes in *Synechocystis*, relevant genes were directly PCR-amplified from the *Synechocystis* genome, fused to the rhamnose-inducible promoter [36] *via* overlap extension PCR, and integrated into a modified variant of the conjugative vector pSHDY containing a nourseothricin resistance instead of spectinomycin, termed pSNDY. The vector backbone also contained the activator *rhaS* from [36]. Lun Yao and Paul Hudson further provided a strain with *gyrB*-targeting sgRNA. For the construction of the additional sgRNA constructs, sgRNA sequences were designed using CHOPCHOP [62], constructed *via* overlap extension PCR and integrated into the vector designed by [37] (Addgene #73224), which inserts into the slr0230 site of the *Synechocystis* genome. Integrative sgRNA plasmids were integrated *via* transformation. Briefly, 10 mL of exponentially grown culture was concentrated to 250 µL, 1 µg–2 µg of pure plasmid was added and the mixture was incubated up to 5 h before plating the entire mixture on BG11 plates. After drying the plates, agar was underlaid with 300 µL of 1 mg mL^−1^ kanamycin stock using a sterile spatula, thereby forming a diffusion gradient. After 1-2 weeks of incubation at 30 *^◦^*C with the lid facing upward, isolated green colonies were carefully transferred to a fresh plate. Over time, positive clones were gradually shifted to higher concentrations of kanamycin (*i.e.*, 4, 8, 12, 20, 40 µg mL^−1^ final concentration in the plate). Complete segregation of mutants was ensured *via* colony PCR. Replicative vectors were introduced into the dCas9 background strain *via* conjugation as described (dx.doi.org/10.17504/protocols.io.ftpbnmn). Clones were selected using nourseothricin (Jena Bioscience, #AB-102L) at a final concentration of 50 µg mL^−1^ and verified *via* colony PCR.

### Culture Conditions

#### Batch Culture Conditions

Plates were freshly streaked before liquid cultivation. In the case of noursethricin, liquid cultures were maintained with 25 µg mL^−1^ while solid media was supplemented with 5050 µg mL^−1^. Spectinomycin and Kanamycin were supplemented at concentrations of 20 µg mL^−1^ and 25 µg mL^−1^, respectively. For pre-culturing and growth experiments, *Synechocystis* strains were cultivated in BG11 medium [63]. Cultivation was performed at 30 *^◦^*C with 150 rpm shaking and continuous illumination of ≈80 µmol m*^−^*^2^ s^−1^ (16 % setting in Infors HT). Aeration was ensured by continuous shaking and CO_2_ enriched air (0.5 %) in an Infors HT multitron chamber. Pre-culturing was performed in 100 mL baffle-free Erlenmeyer shaking flasks with 20 mL cell suspension for three days. After adjusting all different strains to the same OD_750_ 750 ≈ 0.4, growth experiments were performed after one additional day of pre-culturing. For this, 30 mL cultures were incubated in Erlenmeyer shaking flasks for 5 days with a start OD_750_ ≈ 0.25 in biological triplicates.

### Continuous Culture, Online Measurements and Calculations

Continuous culture was performed in a Lambda Photobioreactor and with additional measurement and control devices (Fig. S7) in BG11 medium, supplemented with the required antibiotics, at culture volume *V_£_* = 1 L, aeration with 1 L min^−1^ of CO_2_-enriched (0.5 %) air, agitation by the Lambda fish-tail mixing system at 5 Hz, temperature control at 30 *^%^*C, and pH 8, with 0.5 M NaOH and 0.5 M H_2_SO_4_ as correction solutions. After equilibration to these conditions the reactor was inoculated to a start OD_750_ ≈ 0.5, from 100 mL pre-culture. White light from the Lambda LUMO module was initially increased as a ramp from 42 to 250 photons, then kept constant, and manually decreased to maintain light intensity approximately at ~ 90 µmol m*^−^*^2^ s^−1^ per OD_750_ (Fig. S8F). After the switch to batch culture light was again increased in ramp from 70 to 250 µmol m*^−^*^2^ s^−1^. For evaporation control and continuous culture, the total weight of the reactor setup was kept constant using the built-in Lambda reactor mass control module and automatic addition of fresh culture medium through the feed pump. In batch mode this controls for evaporation of medium. Continuous culture was performed by setting the waste pump to a fixed speed.

#### Calibrations of Measured & Controlled Data

The Lambda Photobioreactor (Fig. S7) was equipped with online monitoring of dissolved O_2_ and pH, and additional monitoring of optical density by a DasGip OD4 module and monitoring of offgas O_2_ and CO_2_ concentrations and the weights of feed and pH control bottles by Arduino-based custom-built data loggers (Fig. S7). The signal from the OD4 probe was calibrated to offline OD_750_ measurements (Fig. S8A-B). For normalizations of glycogen measurements by biomass and for estimation of the biomass density of cells (g_DCW_/mL_cell_, Fig. 5D) a LOESS regression of the OD*_λ_* signal was calibrated to CDW (Fig. S8C-D). The Lambda LUMO light module consistst of a strip of white LEDs and was calibrated to light intensity in µmol m*^−^*^2^ s^−1^ with a Licor light meter (LI-250A) with a spherical sensor bulb (LI-193) (Fig. S8E-F).

#### Calculation of Dilution and Growth Rates

All rates were calculated as slopes of measured data using piecewise linear segmentation [64], implemented in the CRAN R package dpseg (version 0.1.2) [65] (Fig. S9A-D). Growth rates were then calculated as the difference between slopes of measured biomass rate changes (OD*_λ_*, CASY cell counts) and the culture dilution rate corrected for evaporation loss, see Figure S9E-F for details. Cell volume growth rate was calculated as the rate of change of the peaks of the CASY cell volume distributions.

### Biomass & Metabolite Measurements

#### Cell Dry Weight Measurement

To determine the cell dry weight (CDW) 5 mL cell culture was filtered through a pre-dried and pre-weighed cellulose acetate membrane (pore size 0.45 µm) using a filtering flask. After that the membrane was dried at 50 *^◦^*C for 24 h and weighed after cooling. 5 mL of filtered and dried growth medium served as a blank.

#### Optical Density OD_750_ and Absorption Spectra

The optical density (OD_750_) and absorbance spectra were measured on a Specord200 Plus (Jena Bioscience) dual path spectrometer, using BG11 as blank and reference. Samples were appropriately diluted with BG11 before measuring. All topA^OX^ time series samples were diluted 1:4 before recording OD_750_. For absorbance spectra the OD_750_ was adjusted to 0.5 and the absorbances at 750 nm were subtracted from each spectrum.

#### Cell Count and Size Distributions

To determine the cell count 10 µL cyanobacteria culture, pre-diluted for OD_750_ measurement, were dispensed in 10 mL CASYton and measured with a Schaerfe CASY Cell Counter (Modell TTC) using a diameter 45 µm capillary. Cell size was recorded in the diameter range 0 µm–10 µm. Each sample was measured with 400 µL in triplicate runs.

Analysis of the raw data was performed in R. Counted events in the CASY are a mix of live cells, dead cells, cell debris and background signals. Only counts with diameter *d >* 1.5 µm and *d <* 5 µm were considered for the time series experiment (Fig. 5B) while a lower cutoff *d >* 1.25 µm was used for the endpoint measurements (Fig. 2C) to avoid cutting the distribution of the slightly smaller topA^KD^ cells. Since *Synechocystis* cells are spherical, the cell volumes were calculated from the reported cell diameters *d* as 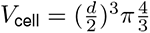.

#### Glycogen Measurement

To determine the glycogen content, 0.5 mL of cell culture was harvested into reaction vessels that had been pre-cooled on ice. After centrifugation at maximum speed for 5 min at 4 *^◦^*C the pellets were flash-frozen in liquid nitrogen and stored at 80 *^◦^*C until further processing. To start the glycogen extraction, the pellet was resuspended in 400 µL KOH (30% w/v) and incubated at 95 *^◦^*C for 2 h. For precipitation, 1200 µL ice cold ethanol was added and the mixture was incubated at 20 *^◦^*C overnight. After centrifugation at 4 *^◦^*C for 10 min at 10 000 g, the pellet was washed once with 70 % ethanol and again with pure ethanol. Afterwards the pellets were dried in a Concentrator Plus speed-vac (Eppendorf) for 20 min at 60 *^◦^*C. Then the pellet was resuspended in 1 mL 100 mM sodium acetate (pH 4.5) supplemented with amyloglucosidase powder (Sigma-Aldrich, 10115) at a final concentration of 35 U*/*mL. For enzymatic digestion samples were incubated at 60 *^◦^*C for 2 h. For the spectrometric glycogen determination the Sucrose/D-Glucose Assay Kit from Megazyme (K-SUCGL) was applied according to the manufacturer’s specifications, but omitting the fructosidase reaction step and scaling down the total reaction volume to 850 µL. Absorbance at 510 nm was measured using a BMG Clariostar photospectrometer.

#### ATP and ADP Measurement

2 mL tubes were preloaded with 250 µL of buffer BI (3 M HClO_2_, 77 mM EDTA). 1 mL culture sample was added, briefly vortexed and incubated on ice for 15 min. 600 µL of BII (1 M KOH, 0.5 M KCl, 0.5 M Tris) were added, vortexed and incubated on ice for 10 min. After centrifugation at 0 *^◦^*C for 10 min at 12 000 g, samples were flash-frozen in liquid nitrogen and stored at 80 *^◦^*C until further processing. Extracts were thawed on ice and centrifuged again at 0 *^◦^*C for 10 min at 12 000 g. 200 µL samples were added either to 320 µL of BIII/PEP (100 mM HEPES, 50 mM MgSO_4_ × 7H_2_O, adjusted to pH 7.4 with NaOH, and 1.6 mM phosphoenolpyruvate (Sigma-Aldrich, 860077) for ATP quantification or BIII/PEP+PK (BIII/PEP with 2 U*/*µL pyruvate kinase (Sigma-Aldrich, P1506) for ATP + ADP quantification, and incubated for 30 min at 37 *^◦^*C. All samples were heat-inactivated at 90 *^◦^*C for 10 min. For luminescence-based quantification, the Invitrogen ATP determination kit was used (ThermoFisher: A22066). 10 µL of each PEP or PEP+PK-treated sample was loaded in a white 96 well plate with solid bottom and kept on ice until the reaction was started. The luciferase master mix was scaled down in volume, and 90 µL of master mix was added to each well. Luminescence was recorded using a BMG Clariostar. ATP concentrations were calculated using a standard curve with commercially available ATP stock solution (Invitrogen).

#### Flow Cytometry and Analysis

Samples were fixed in 4 % para-formaldehyde in 1xPBS (phosphate buffered saline), washed three times in 1xPBS, and stained with the SYTO9 green fluorescent nucleic acid stain from the LIVE/DEAD kit (thermo) according to manufacturer’s instructions. The flow cytometric measurements were taken at the FACS Facility at the Heinrich-Heine University (Dipl.-Biol. Klaus L. Meyer) using a BD FACSAria III. Forward scatter (FSC) and side-scatter (SSC) were recorded. Syto9 was measured with a 530/30 nm filter, and chlorophyll fluorescence was measured with 695/40 nm filter. For each sample 10,000 events (cells, debris and background) were recorded.

Data was exported in `.fcs` format, parsed and analyzed using the `flowCore` R packge [66], and plotted using functions from our in-house `segmenTools` R package.

### DNA and Plasmid Extraction Agarose Gels

#### DNA Extraction

To isolate the DNA, 1 mL culture was centrifuged at maximum speed for 10 min at 4 *^◦^*C, flash-frozen in liquid nitrogen and stored at 80 *^◦^*C. After thawing, the samples were then resuspended in 1 mL 1x TE buffer by pipetting up and down and 100 µL lysozyme (50 mg*/*mL stock solution) was added, inverted, and incubated for 1 h at 37 *^◦^*C. Then 10 µL Proteinase K (20 mg*/*mL stock solution) and 100 µL 20 % SDS were added to the samples and incubated at 37 *^◦^*C for 20 h. The lysed cell suspension was completely transferred to Phasemaker Tubes (ThermoFisher: A33248) and one volume of phenol/chloroform/isoamyl alcohol was added to each tube. The two resulting phases were thoroughly mixed and then separated by centrifugation for 10 min at maximal speed at 4 *^◦^*C and the upper (nucleic acid-containing) phase transferred to a new tube. This was mixed with 100 ng*/*µL RNAse A and incubated for 15 min at 37 *^◦^*C for RNA degradation. Followed by the addition of 1 volume of chloroform/isoamyl alcohol, the centrifugation step was repeated. The upper phase was again transferred and precipitated with 1 volume 2-propanol at 20 *^◦^*C over night. After centrifugation at maximal speed and 4 *^◦^*C for 10 min to pellet the DNA, the supernatant was discarded. The pellet was washed twice with 500 µL ice-cold 70 % EtOH and centrifuged for 10 min at maximal speed at 4 *^◦^*C. After drying the pellets at room temperature, the pellets were dissolved in 30 µL water and the concentration was determined *via* Nanodrop (Thermo Scientific NanoDrop 2000c) .

#### Plasmid Extraction

20 mL of cell culture were mixed with 20 mL of undenatured 99.5 % ethanol, pre-cooled to 80 *^◦^*C, in 50 mL centrifuge tubes and stored at 80 *^◦^*C until processing. After thawing on ice, the supernatant was discarded after centrifugation for 10 min at 4 *^◦^*C and 4000 g. The QIAprep Spin miniprep kit was used for the following steps and adapted and expanded for individual steps. The cell pellet was resuspended in 250 µL Qiagen P1 solution and transferred to 1.5 mL reaction tubes. Then 50 µL lysozyme solution (50 mg mL^−1^) was added, mixed, and incubated for 1 h at 37 *^◦^*C. After the addition of 55 µL of 20 % SDS and 3 µL of proteinase K (20 mg mL^−1^), the reaction mixture was incubated at 37 *^◦^*C for 16 h. Starting with the alkaline lysis with the Qiagen P2 solution, all further steps according to the QIAprep Spin Miniprep Kit were carried out with amounts that were adjusted to the initial volume. Next, the concentrations and quality (260/280, 230/280 ratio) was determined using the Nanodrop (Thermo Scientific NanoDrop 2000c). The T5 exonuclease (NEB: M0363) was used for removal of linear and open circular DNA according to the manufacturer’s protocol and incubated at 37 *^◦^*C for 30 min. The isolated plasmid DNA was further purified with the QIAprep modules. For this, 5 volumes of the PB buffer were added to 1 volume of DNA solution. All subsequent steps were carried out using the QIAprep Spin miniprep kit and the plasmid DNA concentration was determined with the Nanodrop.

### Gel Electrophoresis and Analysis

#### Chloroquine Agarose Gel Electrophoresis

Agarose gels with chloroquine diphosphate (CQ, Sigma: C6628-50G, CAS: 50-63-5) were used to determine the relative migration speed of supercoiled topoisomers. A 1.2 % agarose gel in 0.5x TBE (Roth: 3061.2) was prepared by heating to boiling. After cooling (hand-warm) CQ was added to 20 µg mL^−1^ and the mixture poured into the gel chamber. As running buffer served 0.5x TBE buffer with the same CQ concentration as the gel. For each sample, 120 ng DNA was mixed with loading dye and filled up to 30 µL with water. Next, the voltage source was adjusted 1.8 V cm^−1^. Gels were run for 20 h–24 h covered in foil to protect from light. Next, the gel was washed two times for 30 min in 250 mL 0.5x TBE buffer to remove the CQ. After that, the gel is stained with 25 µL SYBR Gold (ThermoFisher: S11494) in 225 mL 0.5x TBE buffer for 3 h. Then the CQ agarose gel was imaged by a BioRad Imaging System (ChemiDocTM MP). Washing and staining were also performed light-protected. *** This method and its calibration are introduced in more detail in our preprint manuscript at bioRxiv [35]. ***

#### Analysis of Gel Electropherograms

Electropherograms of agarose gels of plasmids were extracted in ImageJ for each lane. Electropherograms from capillary gel electophoresis of total RNA samples (Agilent Bioanalyzer) were parsed from exported XML files using the R package bioanalyzeR (v 0.7.3, obtained from https://github.com/jwfoley/bioanalyzeR) [67]. Electropherograms were then processed in R, using LOESS smoothing and peak detection functions from the msProcess R package (version 1.0.7) (https://cran.r-project.org/web/packages/msProcess/). A baseline was determined in two steps using the msSmoothLoess function . The first step used the full signal and served to determine the coarse positions of peaks. The final baseline was then calculated from the signal after removal of peak values. This baseline was subtracted from the total signal to detect peaks (bands) with the msPeakSimple function from msProcess and calculate peak areas. For the total RNA analysis, the baseline signal stems from mRNA and rRNA degradation fragments, and was used to calculate ratios of rRNA peak areas to the “baseline” area.

### Transcriptome Analysis: Sampling, RNA Extraction and Sequencing

#### RNA Extraction and Processing

Cellular activity was stopped by adding 1 mL culture directly to 250 µL 100 % ethanol supplemented with 5 % phenol, flash-frozen in liquid nitrogen and stored at 80 *^◦^*C until further processing. For RNA extraction, a protocol modified from [68] was used. Briefly, frozen samples were centrifuged at maximum speed at 4 *^◦^*C. After discarding the supernatant, the pellet was resuspended in 1 mL PGTX and incubated at 95 *^◦^*C for 5 min. After cooling on ice for 2 min, 700 µL chloroform:isoamyl alcohol (24:1) were added and the mixture was incubated shaking gently at room temperature for 10 min.The mixture was centrifuged for 10 min at maximal speed at 4 *^◦^*C. The upper phase was transferred to a fresh tube and 1 volume chloroform:isoamyl alcohol was added. After repeating the centrifugation step, the upper phase was again transferred and precipitated with 3 volumes of 99.5 % ethanol and 1/2 volume 7.5 M ammonium acetate at 20 *^◦^*C over night. In cases where low RNA concentrations were to be expected, 1 µL RNA-grade glycogen was added to the precipitation mixture. The RNA was pelleted for 30 min at maximum speed and 4 *^◦^*C, washed twice with 70 % ethanol and resuspended in 30 µL RNase-free water. RNA was DNaseI-digested using commercial DNaseI (ThermoFisher: EN0525), according to the manufacturer’s specifications, but using twice the concentration of reaction buffer. DNaseI-digested RNA was phenol/chloroform extracted again to remove the DNaseI. For precipitation after DNaseI-digest, 1/10 volume of 3 M sodium acetate (pH 5.3), was used instead of ammonium acetate.

#### Quantitative RT-PCR

For qRT-PCR, DNaseI-digested RNA was reverse-transcribed to cDNA using the commercial RevertAid RT (ThermoFisher: K1621) according to the manufacturer’s specifications in a reaction volume of 20 µL. To the 20 µL reaction, 60 µL RNase-free water was added. Of this cDNA dilution, 2 µL per reaction well were loaded, and 8 µL Master Mix prepared from the DyNAmo ColorFlash SYBR Green qPCR-Kit (ThermoFisher: F416L) were added to each 2 µL cDNA reaction. The thermal cycling conditions were as follows: 7 min at 95 *^◦^*C, followed by 40 cycles of 5 s at 95 *^◦^*C and 30 s at 60 *^◦^*C. Data was recorded after each cycle. RT-negative controls and no-template-controls (distilled water) were included for each run. Each sample was loaded in technical triplicates.

Gene expression changes at indicated time points, usually after induction, were then quantified by the ΔΔ*Ct* method [69]:

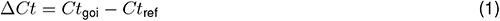

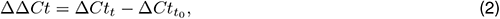

where *Ct*_goi_ and *Ct*_ref_ are the Ct values of a gene of interest (*gyrA*, *gyrB*, *topA*) and of a reference gene (*rpoA*), respectively. The ΔΔ*Ct* value then compares expression at time points *t* after induction with expression at a time point *t*_0_ before induction, and is equivalent to a log_2_ fold change.

#### RNAseq: Total RNA Analysis, Library Generation and Sequencing

RNA quality was evaluated spectrometrically by Trinean Xpose (Gentbrugge, Belgium) and by fragment size distribution on an Agilent 2100 Bioanalyzer with the RNA Nano 6000 kit (Agilent Technologies, Böblingen, Germany). Electropherograms for the endpoint RNAseq samples were exported as XML files for further analysis; see paragraph *Analysis of Gel Electropherograms.* for details.

The Illumina Ribo-Zero Plus rRNA Depletion Kit was used to remove the ribosomal RNA molecules from the isolated total RNA. Removal of rRNA was evaluated with the RNA Pico 6000 kit on the Agilent 2100 Bioanalyzer. RNA was free of detectable rRNA. Preparation of cDNA libraries was performed according to the manufacturer’s instructions for the TruSeq stranded mRNA kit (Illumina, San Diego, CA, United States). Subsequently, each cDNA library was sequenced on an Illumina NextSeq 500 system (2 x 75 nt PE high output v2.5).

#### RNAseq: Read Mapping

The resulting sequence reads were quality trimmed with Trimmomatic v0.33 [70] using standard setting. The quality trimmed reads were subsequently mapped to coding genes of the Synechocystis sp. PCC 6803 reference genome (NC_000911) including the plasmids pCA2._M, pCB2.4_M, pCC5.2_M, pSYSM, pSYSA, pSYSG and pSYSX (CP003270, CP003271, CP003272, NC_005229, NC_005230, NC_005231, NC_005232), and the constructed plasmid pSNDY_Prha_topA-6_119rhaS_20210310 using Bowtie 2 [71].

For the endpoint measurements from batch cultures the log2-fold changes with respect to the controls (EVC) were calculated with the DESeq2 algorithm [45] *via* the ReadXplorer software version 2.0 [72], based on three replicate measurements for each strain (“M-value”). For the time series read-count data were normalized by library sizes to the Transcripts Per kilobase Million (TPM) unit. Missing values at indivdual time points were interpreted as 0 TPM.

### Transcriptome Analysis: Batch and Time-Series Analysis

#### Cluster Analysis

For clustering the time series into co-expressed cohorts, a previously established pipeline was used [73, 74]. Briefly, the time-series of TPM values was arcsinh-transformed:

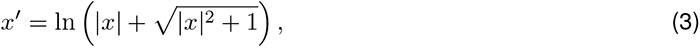

then the Discrete Fourier Transform (DFT) was calculated:

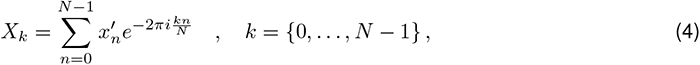

where *x^′^_n_* = {*x*^′^_*0*_,…*x*^′^_*N-1*_} are the (transformed) expression values at time points {*t*_0_*,…, t_N−_*_1_}, and *X_k_*, the DFT, is a vector of complex numbers representing the decomposition of the original time series into a constant (mean) component (at *k* = 0) and a series of harmonic oscillations around this mean. The input time series *x_n_* were RNA-seq samples 2 to 16 (from 0.5 h to 72 h around the time of induction at 0 h), *i.e.*, without the first pre-induction time-point and ignoring the two long-term response samples. Components *k >* 1 were further scaled by the mean of amplitudes at all other components *k >* 1:

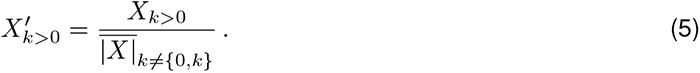

Real and imaginary parts of selected components *X_k=1,…,6_* of the DFT were then clustered with the `flowClust` algorithm [75] over cluster numbers *K* = 2 …10. The clustering with the maximal Bayesian Information Criterion, as reported by `flowClust` (Fig. S12A), was selected for further analyses. Data transformation and clustering were performed by the `processTimeseries` and the `clusterTimeseries2` functions of `segmenTier` and `segmenTools` packages, respectively. The resulting clusters were sorted and colored based on the comparison with diurnal co-expression cohorts (Fig. 6 and S17) for informative plots of the subsequent analyses.

#### Immediate Response Analysis

To estimate the immediate transcriptional response to *topA* overexpression (Fig. 6C, bottom panel) the difference of read counts (TPM) between the means of the two pre-induction time points (−-1 d, -35 min) and the two post-induction time points (5 min, 20 min) was calculated. Transcripts with negative values were labelled as “down”, with positive values as “up”, and 0 or not available values as “nc”.

#### Clustering of Transcription Units

Average expression was calculated for transcription units (TU) reported by ref. [50] from the expression of coding genes they encompass (*via* the “Sense.tags” column of the original data set). The resulting TU time-series was clustered by k-means, using cluster centers from the CDS clustering (Fig. 5) and identical time-series processing. This way protein-coding transcription units could be assigned to the same cluster labels (Fig. S23). Clusters were then sorted by their mean expression peaks, and colored and named according to the diurnal time of their peaks.

#### Clustering of Diurnal Transcriptome Data

Diurnal expression data from [48] were obtained from GEO (GSE79714) and genes summarized as the mean over all associated probes. These expression values were clustered the same was as described for the RNA-seq data: all time-series were DFT-transformed and amplitude-scaled DFT components 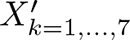 (eq. 4–5) were clustered with segmenTier functions `processTimeseries` and `flowclusterTimeseries`, using the maximal BIC clustering with *K* = 5 clusters.

#### Cluster-Cluster Overlap Tests

Categorical enrichments, *e.g.*, coding gene co-expression cohorts vs. gene annotations, were analyzed by cumulative hypergeometric distribution tests (R’s phyper) using `segmenTools’s clusterCluster` function and the `clusterAnnotation` wrapper for GO and and protein complex analysis, which compares overlaps of each pair of two distinct classifications into multiple classes, and stores overlap counts and p-values (enrichment tables) for informative plots (see “Enrichment Profile Plots”).

#### Enrichment Table Sorting

For intuitively informative plots the enrichment tables were sorted. Table rows were sorted along the other dimension (table columns) such that all categories enriched above a certain threshold *p*_sort_ in the first column cluster are moved to the top, and, within, sorted by increasing p-values. Next, the same sorting is applied to all remaining row clusters for the second column cluster, and so on until the last column cluster. Remaining row clusters are either plotted unsorted below a red line or removed. This is especially useful to visualize enrichment of functional categories along the temporal program of co-expression cohorts, *e.g.*, Figure 6B. This sorting is implemented in segmenTools’ function `sortOverlaps`.

#### Cluster t-Test Profiles

To compare clusters (co-expression cohorts) with numerical data, here the *log*_2_ fold-changes of transcript abundances in induced gyr^kd^ and topA^OX^ strains, we developed the `segmenTools’` function `clusterProfile`. For each cluster a two-sided `t-test` was performed (R base function `t.test`, incl. Welch approximation for different sample sizes), comparing the distribution of values of the cluster with all other values. The reported *t* statistic and the *p*-value were stored for each test. The resulting t-test profile was stored for informative plots (see “Enrichment Profile Plots”).

#### Enrichment Profile Plots

The results of cluster enrichment tests (cluster-cluster overlap tests and t-test profiles) were visualized as colored table plots (*e.g.* Fig. 6B, C), using `segmenTools’` function `plotOverlaps`. For the categorical overlap tests, the total counts of overlapping pairs are plotted as text, and for t-test profils the rounded *t* statistic. The text color is black or white based on a p-value cutoff *p*_txt_ (as indicated).

The field background colors scale with log_2_(*p*) of the reported p-values, where the full color corresponds to a minimal p-value *p*_min_ cutoff (as indicated). For categorical enrichment tests the full color is black and other colors are selected from a gradient to white. For numerical tests, the sign of the *t* statistic is used to determine a color to indicate the direction of change: red for negative values (downregulated) and blue for positive values (upregulated).

#### Motif Scan and DNA Structure Analysis

The genome was scanned for short sequence motifs using custom-built R code based on the `str_locate_all` function of the `stringr R` package [76] into a vector of 0 and 1 for each genome position, where 1 indicates occurence of the motif under consideration. Motif occurence vectors upstream and downstream of start codons or transcription start sites were extracted from the genome vector and aligned into a matrix (columns: positions around the alignment anchor, rows: all genomic sites under consideration). The occurence of a motif in all sequences of a cluster were counted at each position *i* in 66 bp windows surrounding the position. Cumulative hypergeometric distribution tests (R’s phyper) were performed to analyze statistical enrichment or deprivation of a given motif within the 66 bp window of all genes in a cluster vs. the same window in all genes in the total analyzed set (all clustered and aligned sequences), and the p-values *p_i_* at position *i* stored. In the resulting plots of the mean position-wise motif occurence the size of the plotted data point at position *i* was scaled by the enrichment or deprivation p-values to emphasize regions of significant difference (Fig. 6D). The maximal size was determined by the minimum p-value in each test series (Figure panels). The point style (closed or open circles) indicates the directionality of the test (enriched or deprived). The significance points are only shown at every third or tenth position to avoid overlaps.

#### Other Data Sources

Genome sequences and annotation were downloaded from NCBI, see “RNAseq: Read Mapping” for the RefSeq IDs. The gene “categories” annotation was downloaded on 2017-09-23 from CyanoBase [77]: http://genome.annotation.jp/cyanobase/Synechocystis/genes/category. txt. Gene Ontology annotation was downloaded from the UniProt database (2021-03-20, organism:1111708) [78]. Datasets from other publications were all obtained from the supplemental materials of the indicated publications.

---

**Figure S1.**
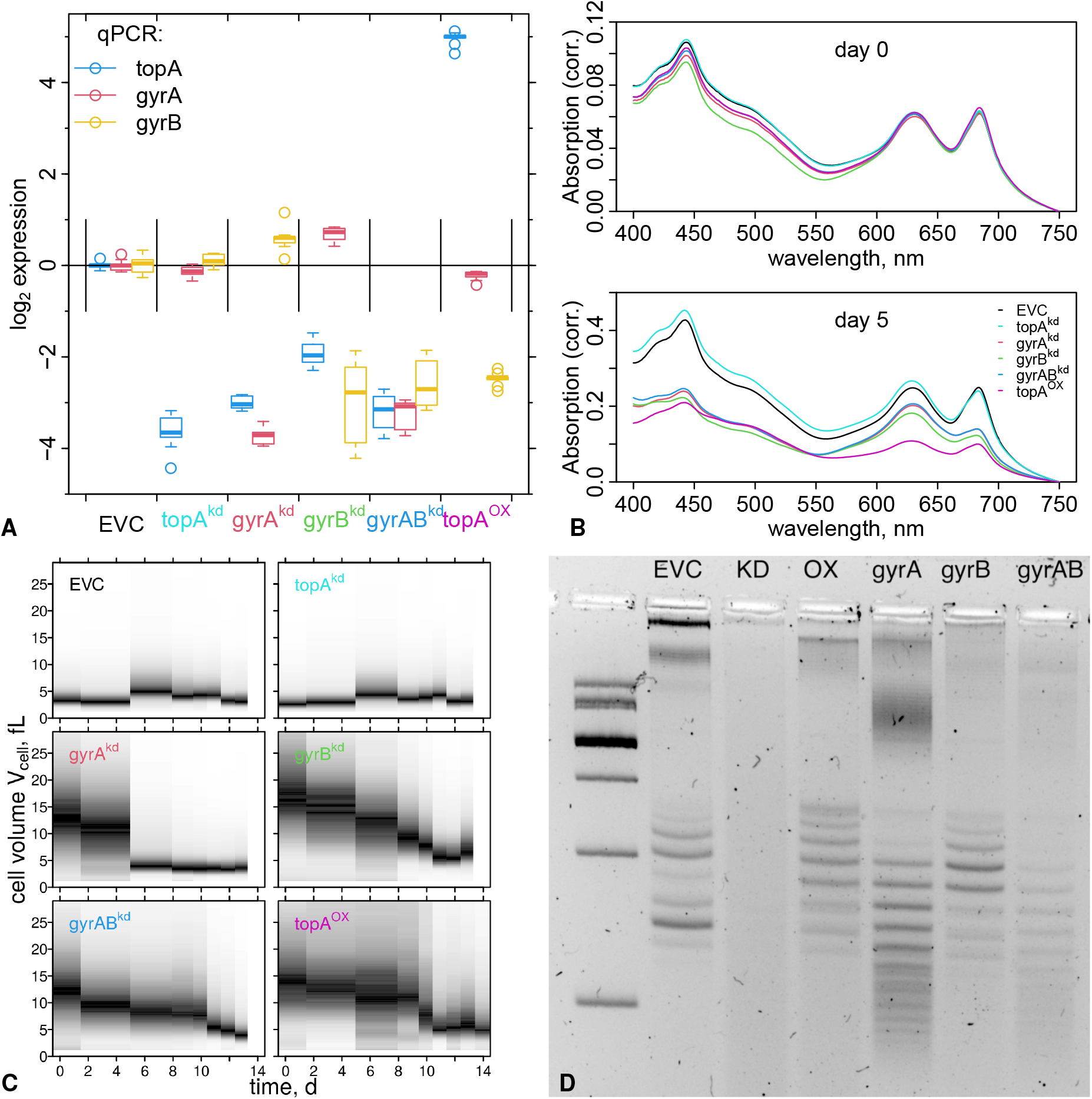
Batch Culture Endpoint Measurements. See Figure 2 for details. **A:** RT-qPCR results using *rpoA* as reference gene. Boxplots of 9 technical replicates (3 samples, each measured 3x) . **B:** Absorption spectra at incolution and harvest times. **C:** Recovery without inducer: all cultures were re-inoculated in fresh BG11 with antibiotics but without the inducers (aTc and rhamnose) and the cell size distributions measured at indicated time points. **D:** Chloroquine-agarose gels (1.2% agarose, 0.5x TBE and 20 µg mL^−1^ CQ) of plasmids extracted at harvest time (5 d).

**Figure S2.**
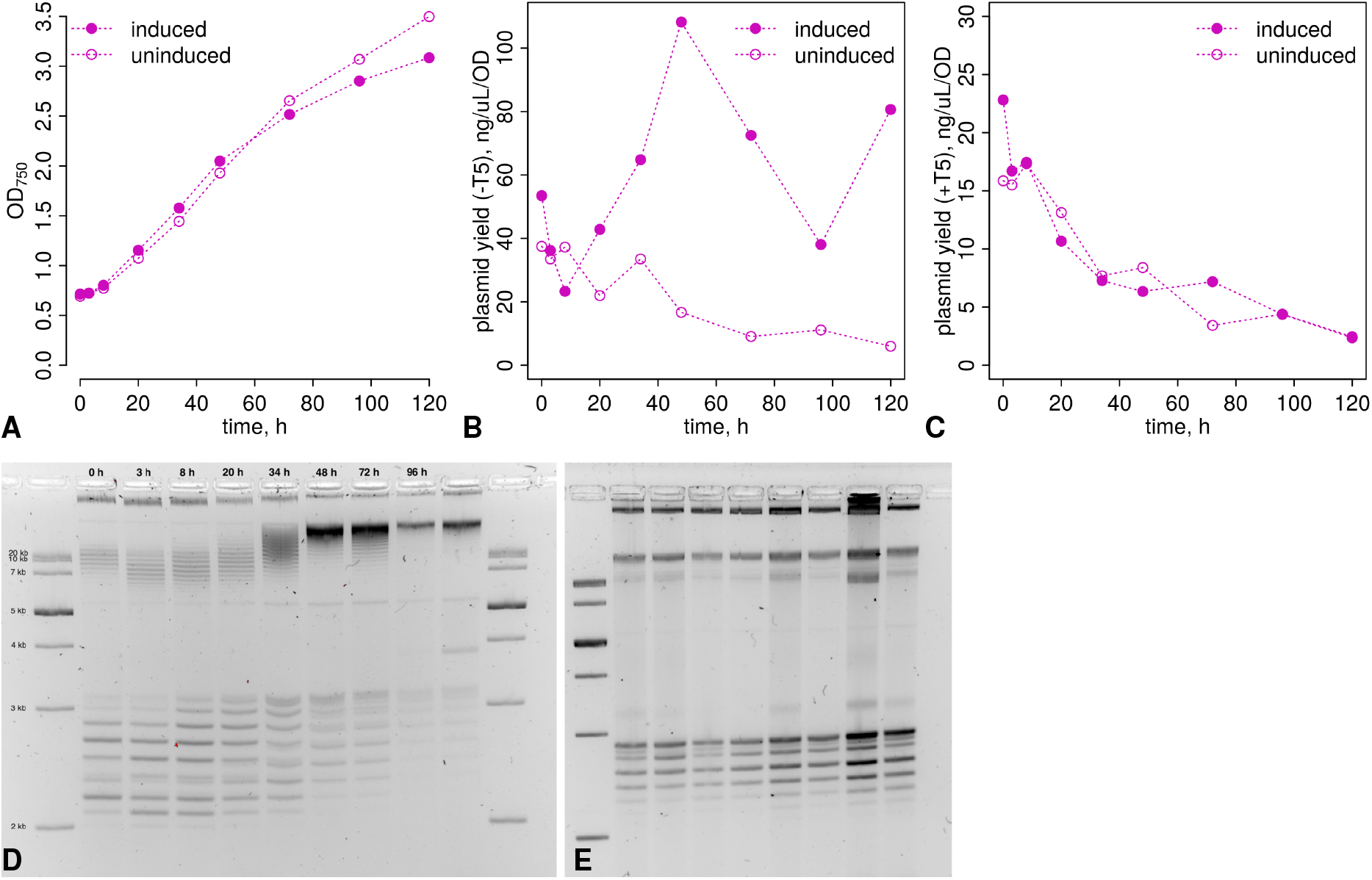
Plasmid Supercoiling Gels. **A:** Growth curves of topA^OX^ strain, induced with 1 mM rhamnose at time 0 h and uninduced control. A starter culture was split into 8 cultures at 0 h, each harvested at the indicated time points for OD_750_ measurement and plasmid extraction. **B & C**: Yields of plasmid extraction over time, each normalized to the OD_750_ (A), and before (B) and after (C) treatment with the T5 exonuclease to remove all non closed circular DNA. **D & E**: Chloroquine-agarose gels (1.2% agarose, 0.5x TBE and 20 µg mL^−1^ CQ) of plasmids extracted from topA^OX^ strain cultures (A), induced (B) and uninduced (C). Note that only the gel of the induced culture (B) was run for a longer time (XYZ) to get a better separation of topoisomers of the pCC5.5_M plasmid. Samples are ordered by sampling times (see A) from left to right on both gels. The 120 h sample is missing in (E).

**Figure S3.**
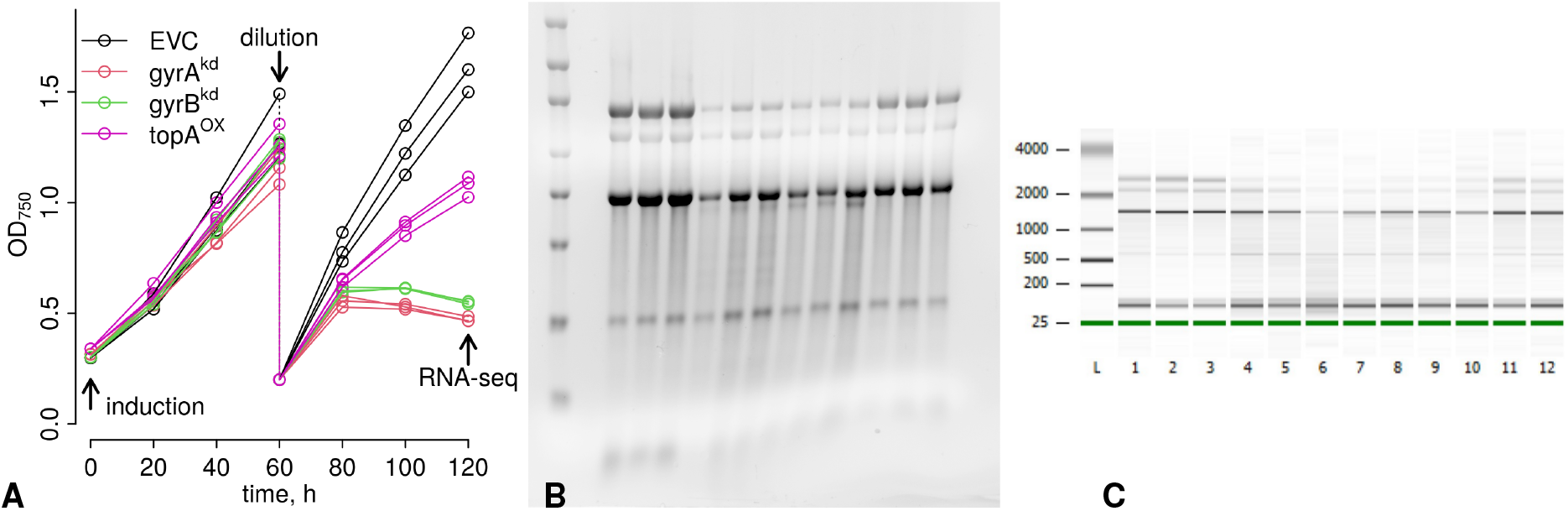
Growth Curves & RNA Extraction for RNA-seq Experiment. **A:** Growth curves of triplicate cultures (split upon induction at 0 d).**B & C:** Total RNA compositions were analysed by a formaldehyde-agarose gel (B, 500 ng RNA per well) and by capillary gel electrophoresis (Agilent Bioanalyzer); electropherograms of these samples were analyzed for Figures 4, S5 and S6. Sample lane order are identical in (B) and (C) and comprise of triplicates from EVC (lanes 1-3), gyrA^kd^ (lanes 4-6), gyrB^kd^ (lanes 7-9) and topA^OX^ (lanes 10-12).

**Figure S4.**
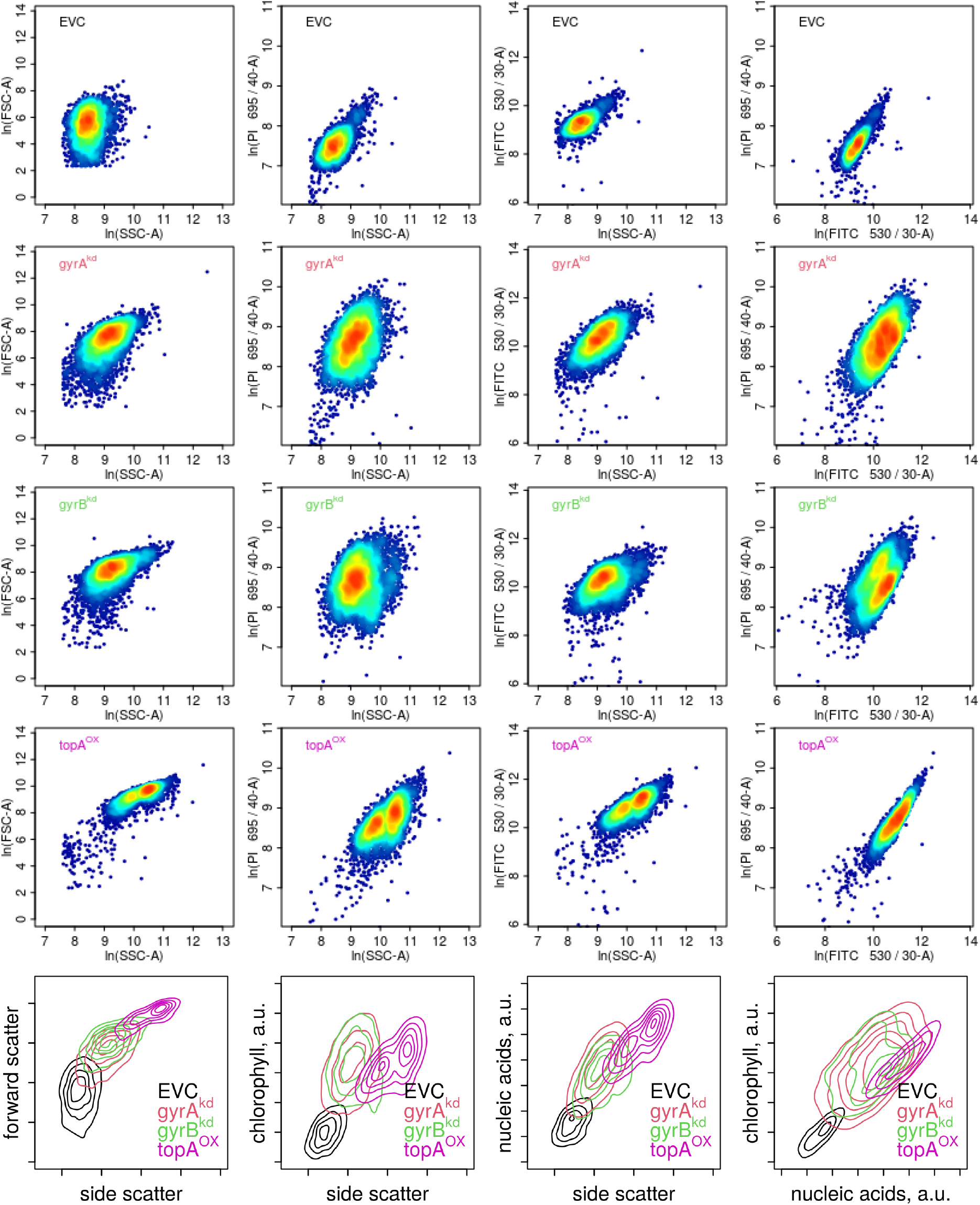
Flow Cytometry. Cells from the cultures used for RNA-seq and total RNA analysis (Fig. 4) were fixed in para-formaldehyde and stained with Syto-9, a nucleic acid fluorescence marker, and analyzed by flow cytometry. The data was gated by the side scatter signal (SSC-A*>* 2000) and the forward scatter signal (FSC-A*>* 10) to filter debris and background signals. Forward scatter (FSC-A) is proportional to cell size, side scatter (SSC-A) reflects cytoplasmic granularity and morphology; the FITC fluorescence channel (530/30 nm) excites the Syto-9 stain, and the PI channel (695/40 nm) excites chlorophyll. The natural logarithm (ln) of all data was plotted. Colors reflect local density (red: high, blue: low). The bottom panels show a zoom into the data, comparing the two strains with the most extreme values, EVC and topA^OX^ (white: high local density).

**Figure S5.**
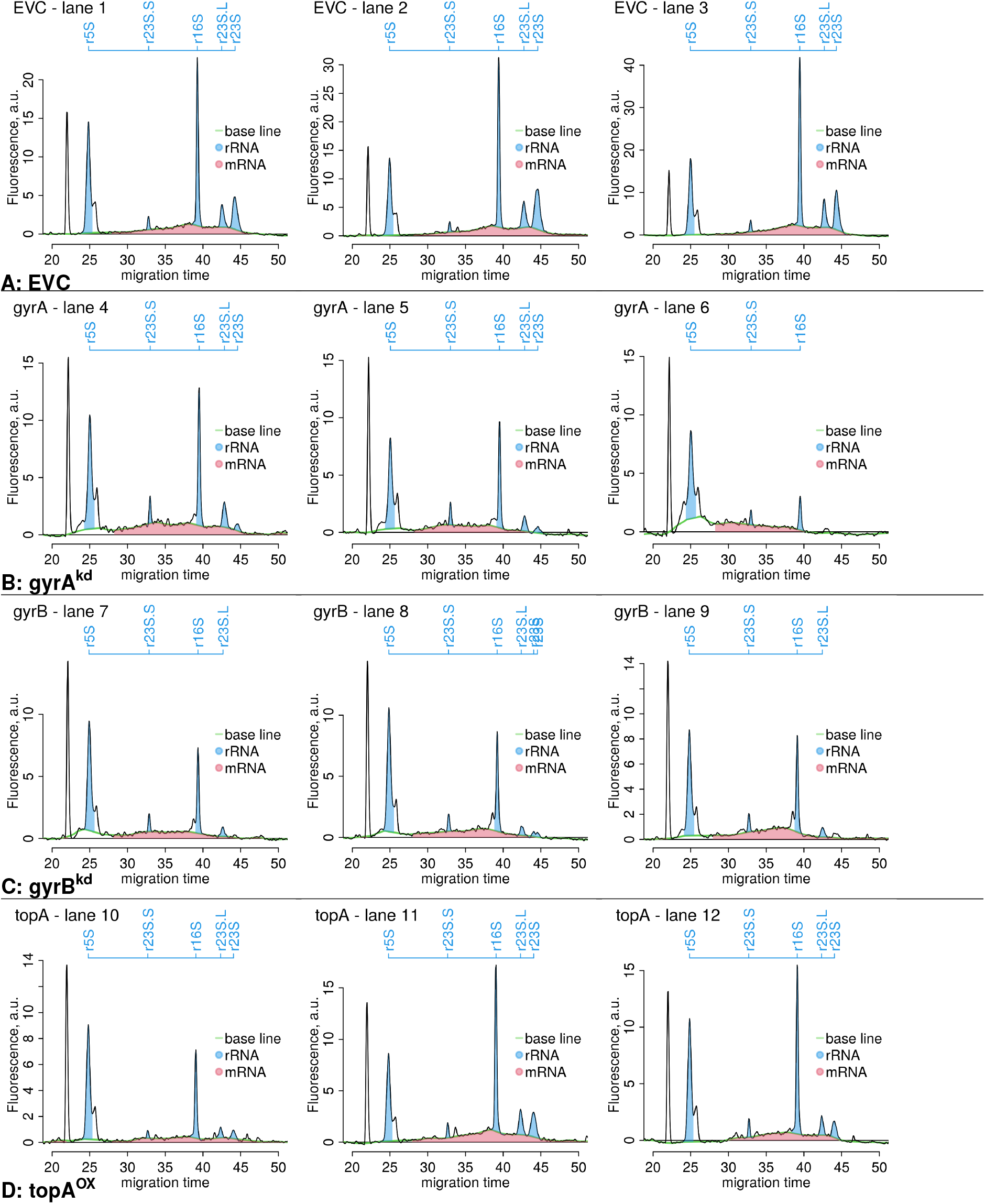
Total RNA Electropherogram Analysis. Electropherograms of the capillary gel electrophoresis (Fig. 4A), exported as XML files from the 2100 Bioanalyzer software. **A–D:** are each the triplicate samples for the indicated strain. Samples are the same as shown on the formaldehyde-agarose gel in Figure S3B and subsequently used for RNAseq analysis (Fig. 4B,C). Data was parsed into R with `bioanalyzeR` (v 0.7.3) [67]. Baseline and peak detection and quantification were performed in R, see Methods paragraph *Analysis of Gel Electropherograms* for details. The areas of the blue peaks were further analyzed as relative rRNA species abundances (see top axis annotation). r23S.S and r23S.L are short and long fragments of the 23S rRNA typically seen in *Synechocystis*, the other peaks were assigned to 5S, 16S and (full length) 23S rRNA. The red area under the baseline (green) and *>* 200 bp was used as relative mRNA abundance for the ratios in Figure 4B,C and S6. The 200 nt length cut-off was based on the size calibration to the RNA ladder lane as reported by the `bioanalyzeR` parser. The peak on the left is the 25 nt lower size marker.

**Figure S6.**
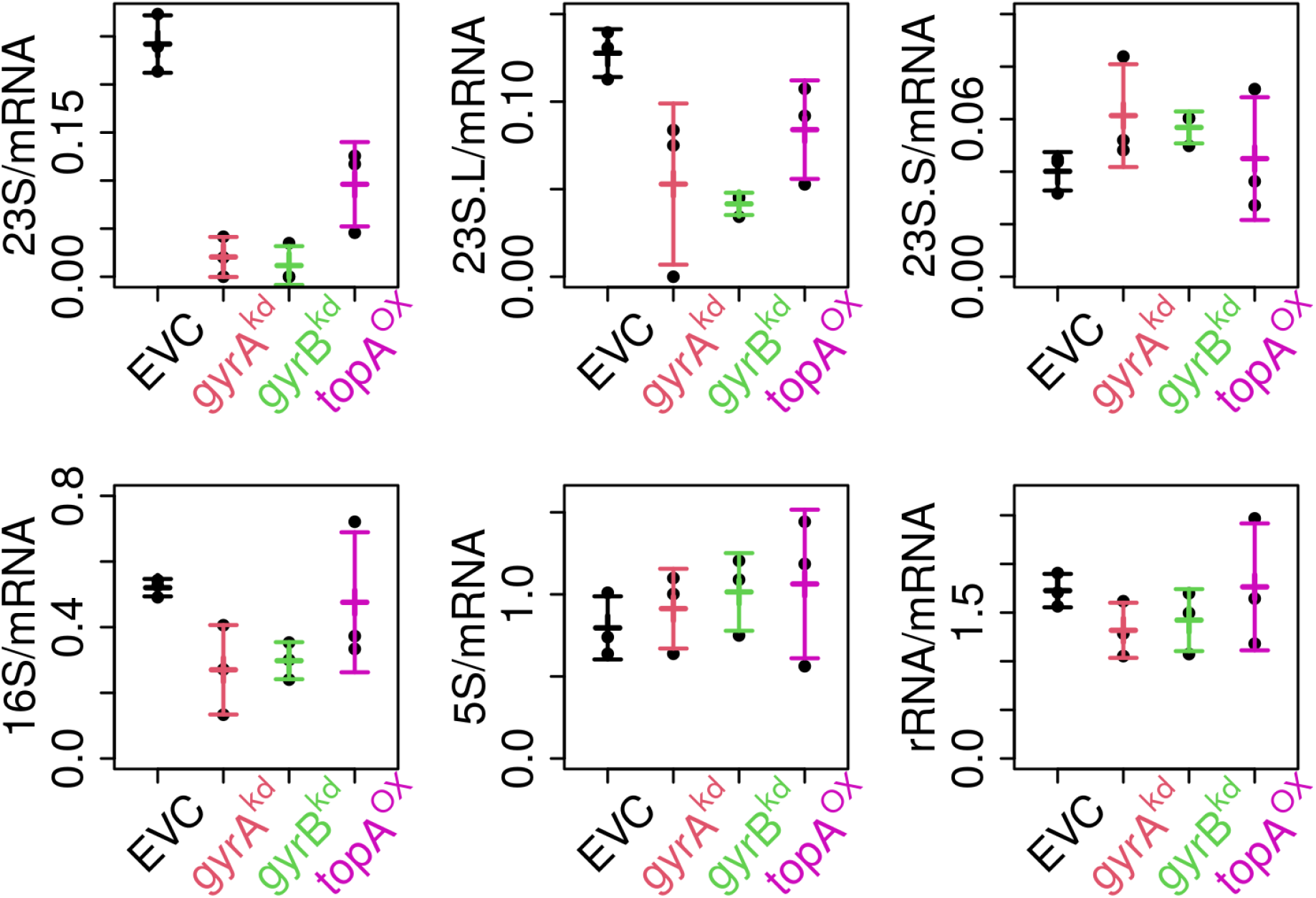
rRNA vs. mRNA: Relative Abundances. Ratios of the indicated rRNA peak areas to the baseline (mRNA) area, the blue and red areas in Figure S5, respectively. The error bars indicate the mean and standard deviation of the three replicates.

**Figure S7.**
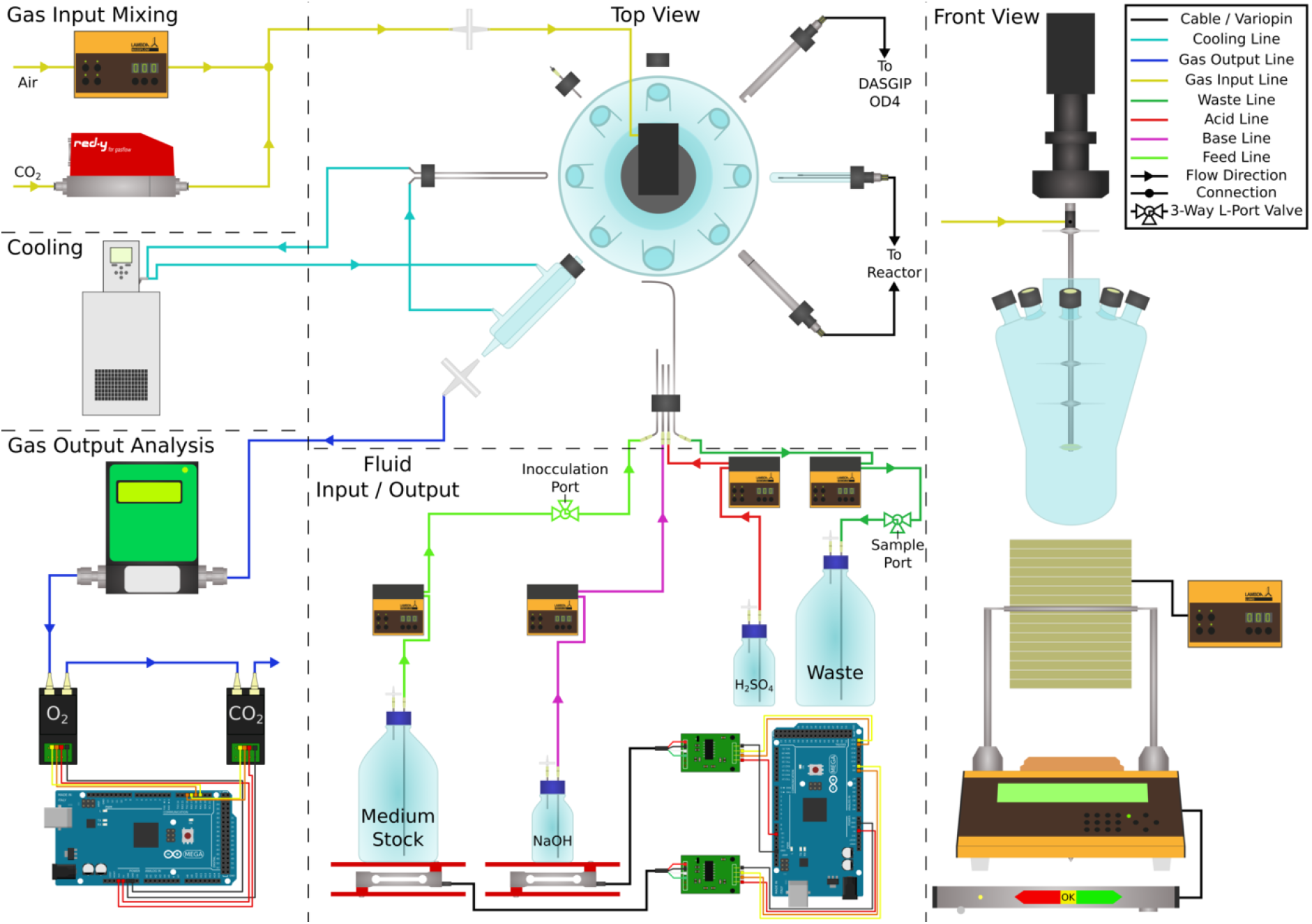
Photobioreactor Setup. A schematic overview of the cultivation setup showing the Lambda minifor bioreactor in front and top-down views alongside its external components and custom expansions. The gas input mixture is generated by a Lambda MASSFLOW 5000 gas flow controller and a Voegtlin red-y smart controller, which regulate the flow of compressed air and CO_2_ respectively. This input gas mixture is then introduced into the cultivation vessel via the sparger at the end of the agitation unit. The offgas condenser as well as the reactor’s cooling finger are part of a water cooling circuit which is regulated by a Lauda Eco Silver thermostat set to 16 *^◦^*C. An Aalborg Massflow Meter monitors the flow rate of the culture’s offgas before it is lead through a custom microcontroller-based gas sensor array in order to evaluate its O_2_ and CO_2_ content. The reactor actively regulates the culture’s pH and temperature values by controlling its heating compartment as well as the Lambda Preciflow peristaltic pumps which are attached to NaOH and H_2_SO_4_ stock bottles, each 0.5 M. Additional culture parameters are monitored by a dissolved O_2_ probe attached to the reactor and an OD4 probe connected to a DASGIP OD4 device. An additional set of peristaltic pumps is attached to the culture’s medium stock and waste containers in order to control the reactor’s volume and medium turnover. The reactor weighting module enables the system to operate under chemostatic conditions. This is achieved by manually configuring the medium feed peristaltic pump at a constant speed in order to achieve a desired medium turnover rate while automatically regulating the waste pump speed to keep the total reactor weight constant. Additionally, a custom microcontroller-based scale setup is monitoring the weight of both the medium and NaOH stock bottles, which allows for the calculation of medium and base pump rates from the recorded data. The culture’s illumination is provided by the Lambda LUMO modules, an LED strip fitted around the cultivation vessel.

**Figure S8.**
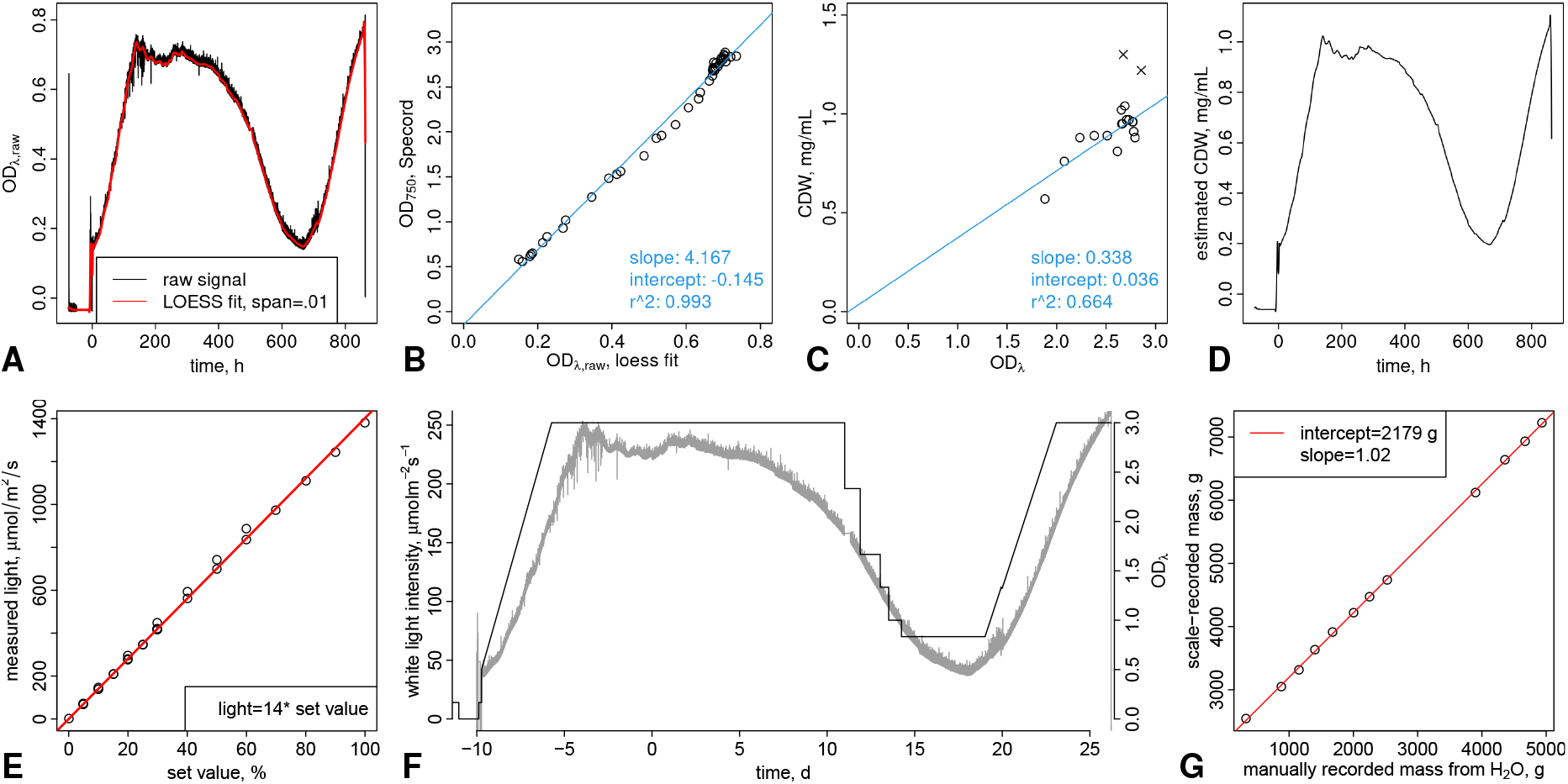
Calibrations. **A:** LOESS regression (R loess) of the raw signal (resolution ca. 1 sec) from the DasGip OD4 module (OD*_λ,_*_raw_). **B:** calibration of the LOESS fit of the OD*_λ,_*_raw_ signal to offline OD_750 nm_ by linear regression (R *lm*). The calibrated signal is used throughout the document and denoted OD*_λ_*. **C:** calibration cell dry weight (CDW) to the OD*_λ_* signal. Data points marked by X were removed as outliers. **D:** the LOESS fit of the OD4 signal was then used to estimate CDW for all time points. **E:** calibration of the Lambda LUMO light module with a Licor light meter (LI-250A) with a spherical sensor bulb (LI-193). **F:** time-series of set and calibrated light intensities (black line, left y-axis) compared to the OD*_λ_* time-series (gray line, right axis). The light intensity was manually adjusted to avoid high-light stress in the culture during biomass decrease. **G:** The Arduino-based scales where calibrated prior to the experiment (not shown). During the experiment the liquid level on the 5L feed bottle was marked regularly, and the mass of water filled to these marked was recorded on a benchtop scale (Kern XZY) after the experiment to test consistent performance. The recorded mass was reproduced sufficiently well (red line: linear regression): the intercept of the linear regression corresponds to the mass of the empty feed bottle and the slope was ≈ 1. Since the manual marks on the bottle are more error prone than the pre-calibration, we did not re-calibrate the data but relied on the recorded mass for calculation of the dilution rate.

**Figure S9.**
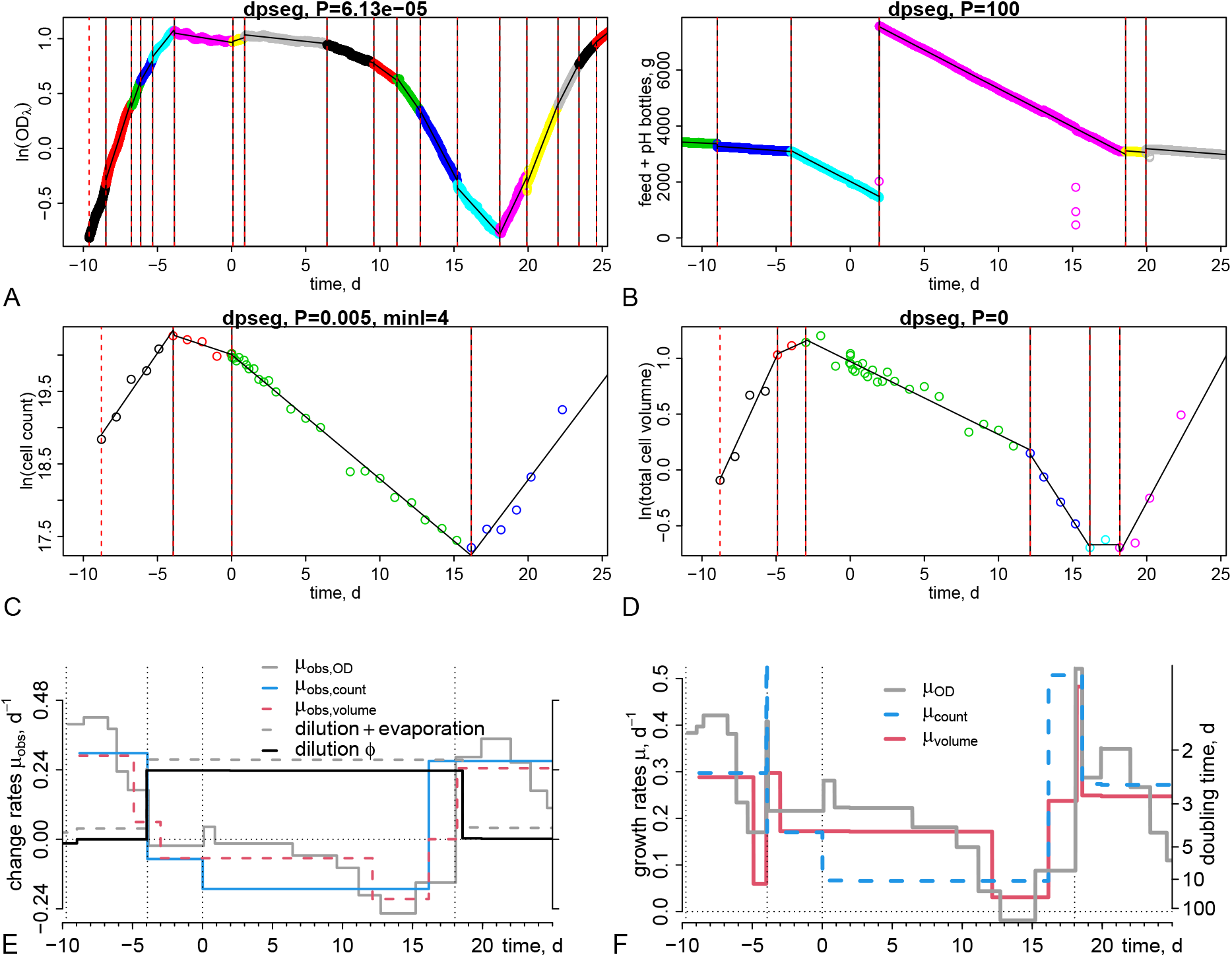
Calculation of Dilution and Growth Rates. All rates were calculated from the slopes of measured data (or of their natural logarithms as indicated) using piecewise linear segmentation with the R package dpseg. The plots in **A-D** were generated by dpseg and the vertical lines indicate borders of the piecewise segments, and the used penality parameter *P* is shown in the plot title on the top axis. The minimal segment length paramater minl was only used in (C). **A**: the calibrated OD*_λ_* signal (1 sec resolution) was smoothed with a moving average and window size 15 and interpolated at 300 sec intervals. **B**: sum of the recorded weights of edium feed and pH control bottle weight; outliers (faulty measurements or bottle changes) were removed and data interpolated at 300 sec intervals. **C:** the total cell count for each CASY measurement, single measurements and means of technical duplicates. **D:** the total cell volume, calculated as the integral of the single cell volume distribution, for each CASY measurement, single measurements and means of technical duplicates. **E:** Observed rates. The (negative) slopes of the summed bottle weight changes (B) reflect the amounts added to the reactor culture by the Lambda reactor mass control system, assuming 1 g/mL density. The total culture dilution rate (dashed gray line, “dilution + evaporation”) is obtained by division by the culture volume (*V_£_* = 1 L). The liquid loss by evaporation is seen at times before onset of continuous culture (time -4 d) and is subtracted to obtain the actual dilution rate *φ* (black line). The slopes of the change of the natural logarithms of the OD*_λ_* signal (A), the total integrated cell volume (B), and the cell counts (C) are the observed change rates *µ*_obs,OD_ (gray line), *µ*_obs,volume_ (red line) and *µ*_obs,count_ (blue line), respectively. **F:** The culture growth rates *µ*_OD_ (gray line) and *µ*_count_ (blue line) and *µ*_volume_ (red line) were calculated as the difference between observed change rates and the culture dilution rate: *µ* = *µ*_obs_ *− φ*.

**Figure S10.**
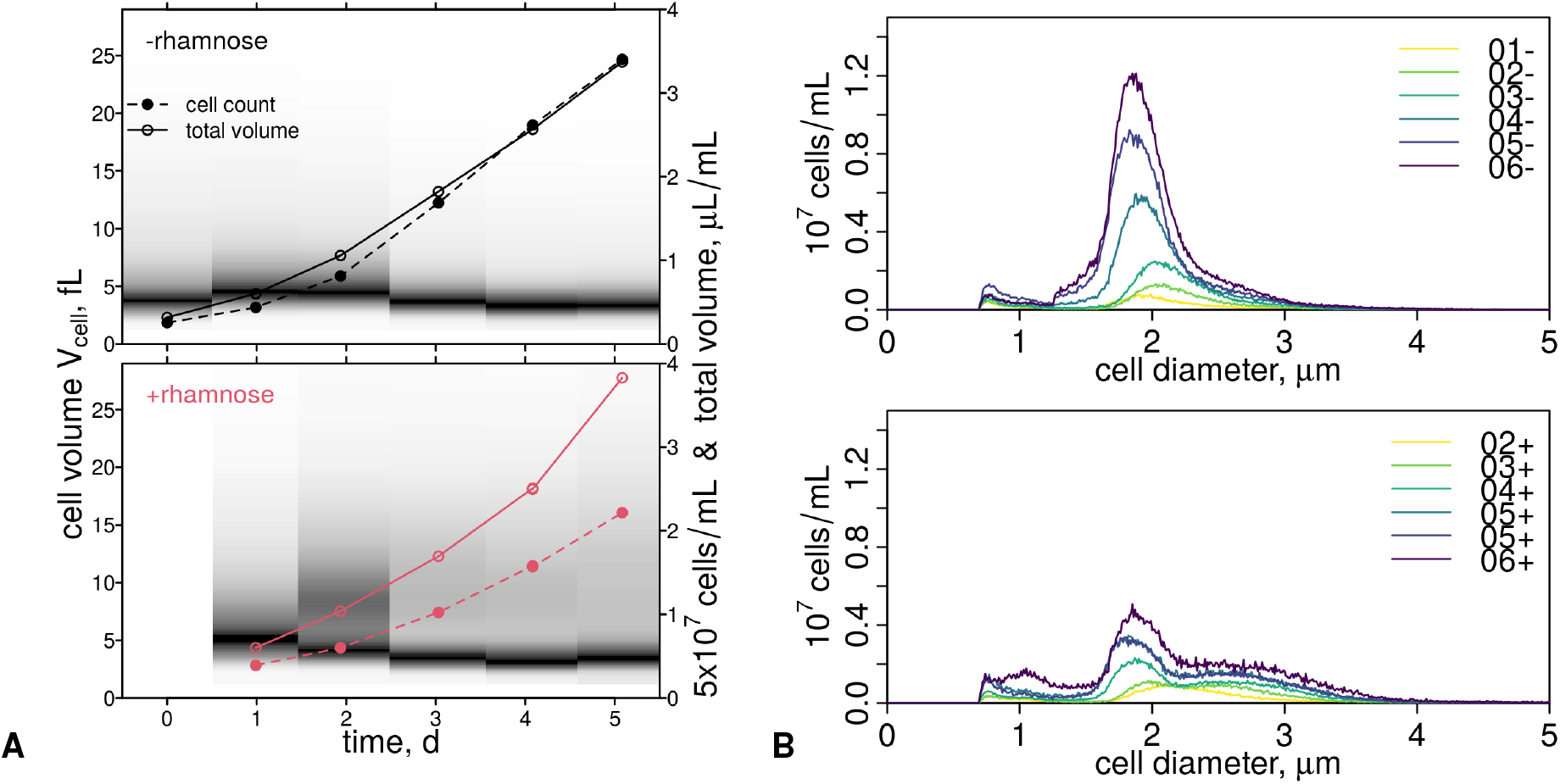
Re-Induction of Bioreactor Time-Series Cultures. After harvest, cells from the bioreactor timeseries experiment (strain topA^OX^) were spinned down, washed and re-inoculated in fresh BG11 with antibiotics and without (top panels,-rhamnose) or with (bottom panels, +rhamnose) in shake flasks (batch culture), and cell count and volume distribution measured daily with the CASY cell counter. **A:** cell volume distributions as gray-scale (left axis) and cell count and total cell volume with open and closed points (right y-axis). **B:** Distributions of the cell diameter as reported by the CASY cell counter and from which cell volumes were calculated. Cells grew normally without further topA^OX^ induction and showed the initial transient volume increase that was observed in all experiments without induced volume growth (*e.g.* EVC and topA^KD^ in Fig. 2B and Fig. S1C). After re-induction of topA^OX^, only a fraction of cells again showed in increase in cell volume.

**Figure S11.**
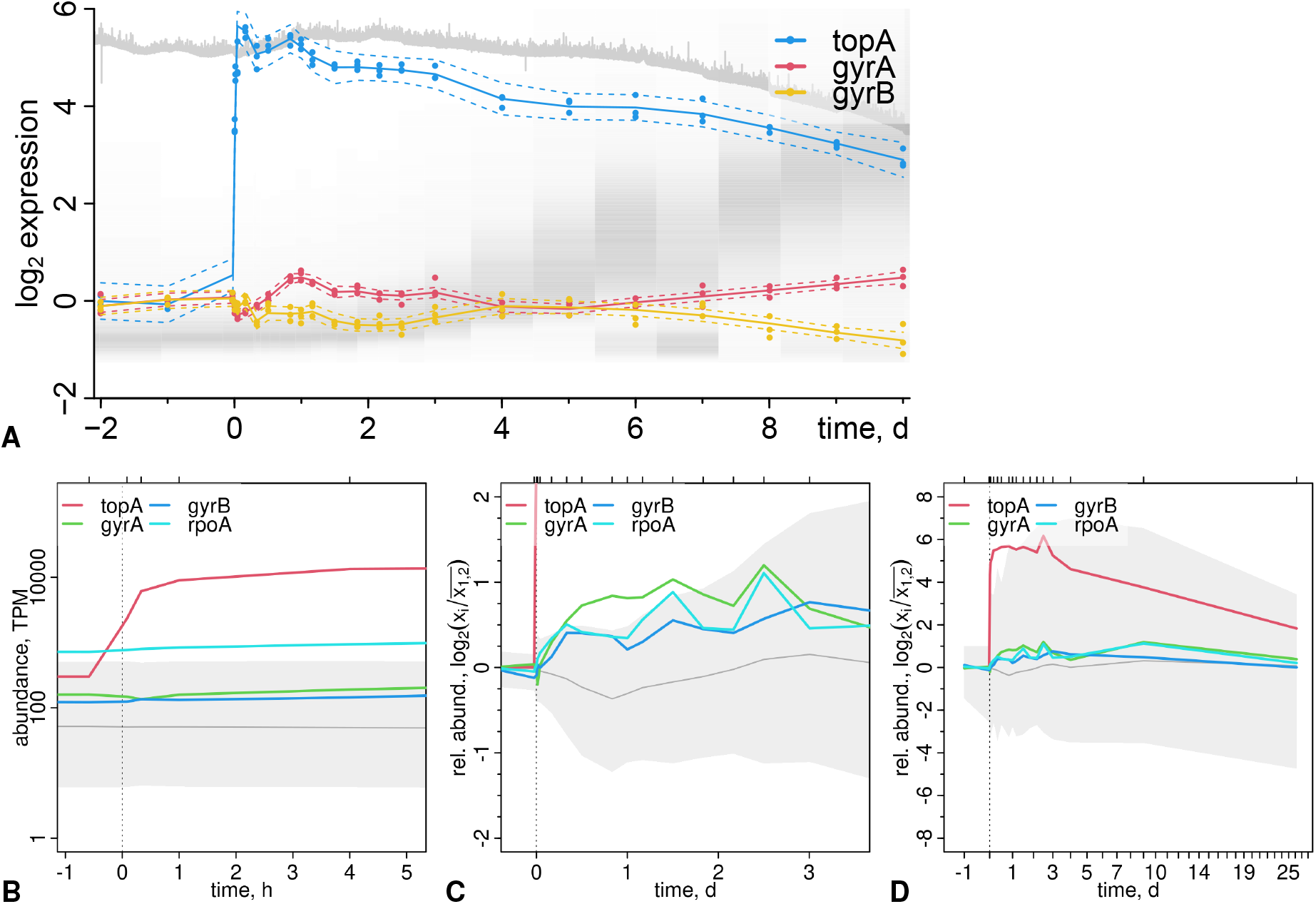
RT-qPCR vs. RNA-seq. **A:** The topA, gyrA and gyrB genes were also measured by RT-qPCR using *rpoA* as a reference “house-keeping” gene. **B-D:** The tested genes and the house-keeping gene in the RNA-seq data at different x-axis zoom levels and for raw TPM read-count data in (B) and log2-fold change over the mean of the two pre-induction samples (C, D). Notably, *rpoA* expression in RNA-seq data increases with a periodic pattern and this should have affected the RT-qPCR measurements. Considering this, the RT-qPCR and RNA-seq data are consistent: gyrA initially decreases and gyrB increases (*<* 1 h); later (*<* 3 d) gyrA increases more (than *rpoA*) and gyrB less (than *rpoA*); followed by a phase of roughly equal expression (4-5 d) and again bifurcation of expression values (*>* 6 d).

**Figure S12.**
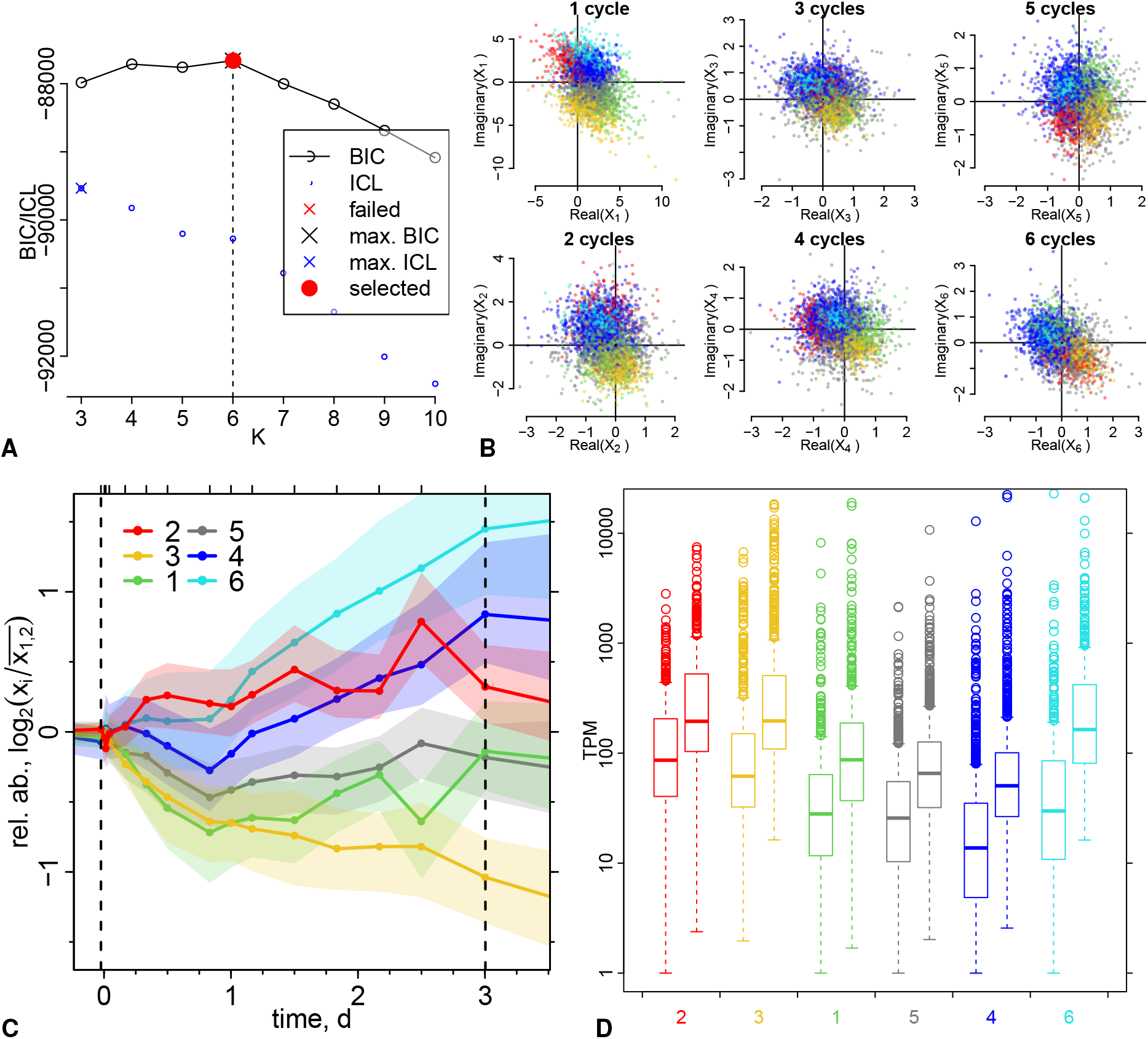
Clustering & Total Read-Count Distribution. A: Bayesian Information Content (BIC) as reported by `flowClust` for clustering of selected scaled components 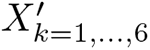 of the Discrete Fourier Transform (DFT, eq. 5)) of the raw TPM data over varying number of cluster centers (*K*). The maximal BIC was reached for a classification into *K* = 6 distinct clusters (co-expression cohorts). This clustering was chosen for further analysis. **B:** real and imaginary parts of the DFT that were used for clustering (R package flowClust [75]). Colors already indicate the final cluster assignments of each transcript at *K* = 6 (A). **C:** Cluster medians (solid lines) of the relative transcript abundances (rel. abund.). For each transcript the log_2_ of the ratio of read-counts at time points *i* to mean of the two samples before induction (*i* = 1, 2, at –1 d and –1 h) was calculated (points indicate the sampling time points *i*). The transparent ranges indicate the 25% and 75% quantiles of each cluster. Only the time points within to two vertical lines were used for clustering. **D:** Cluster-wise distributions (boxplots) of minimal (left) and maximal (right) read-count values (TPM) of each transcripts.

**Figure S13.**
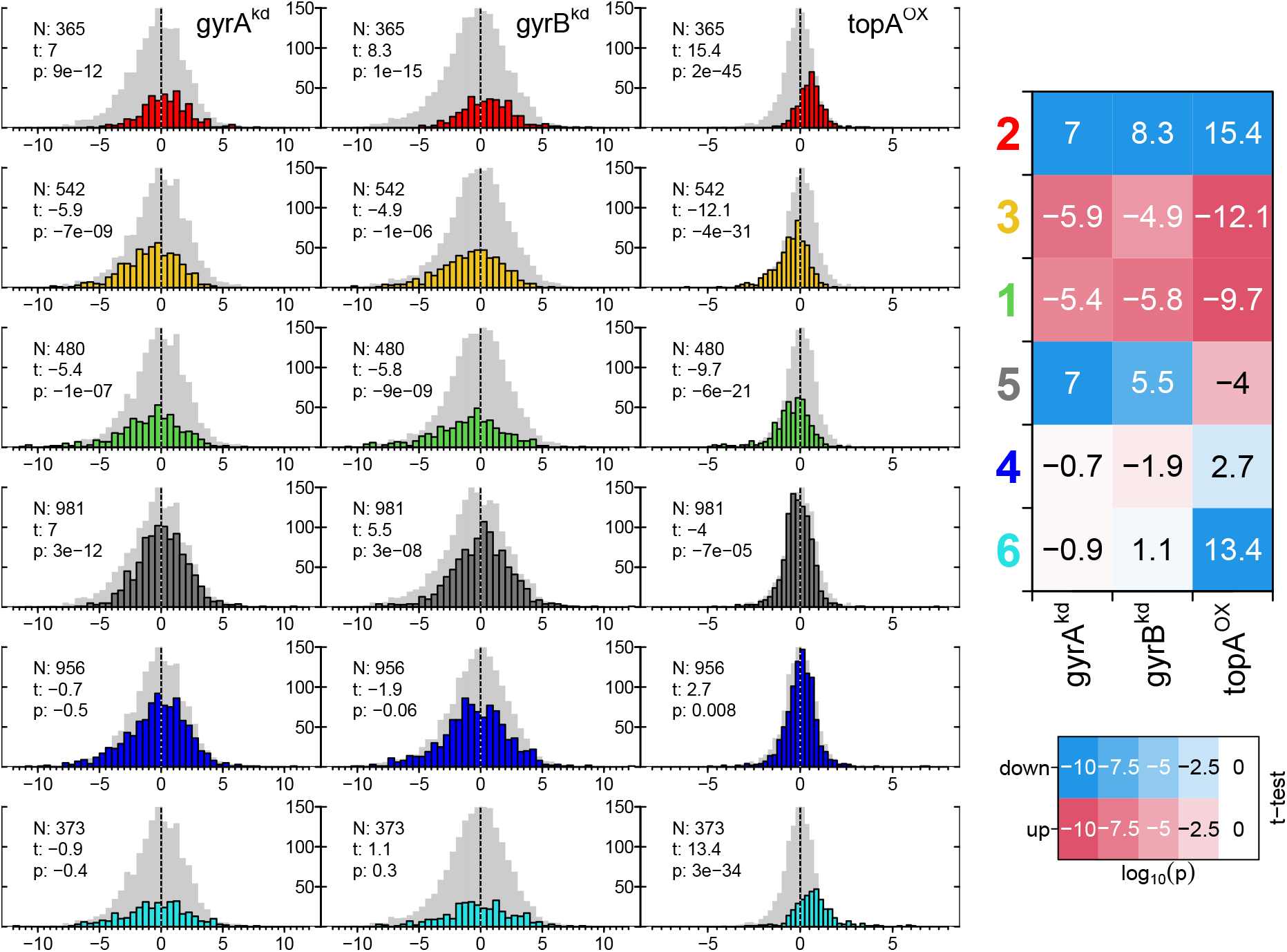
Co-Expression Cohorts in the Endpoint RNA-seq and Construction of t-Test Profiles. **A:** Distributions of the log_2_ fold-change of transcript abundances in the three strain endpoint experiment for each of the co-expression clusters derived from the topA^OX^ time series data. The gray background shows the distribution of all other transcripts. The y-axis are the counts for the colored distributions, while the gray background distributions are densities (without axis). For each cluster a t-test was performed (base R function t.test) against all transcripts not in the cluster, and the cluster sizes *n*, and the *t*-values and the *p*-values from each test are shown in each plot. The total number of transcripts with expression values were 3676 for gyrA^kd^ and gyrB^kd^, and 3680 for topA^OX^. **B:** A t-test profile plot is constructed from the t-test results in (A). A negative *t*-value indicates that the tested cluster transcripts have a lower mean abundance than all other transcripts and this is indicated by a red color field, the rounded t-value is shown in the fields; blue indicates a positive *t*-value and higher mean abundance. The *p*-value is converted to a transparency value for the red and blue colors (along a color palette from red/blue to white), such that the full color is reached for *p ≤ p*_min_, and for higher p-values the transparency scales with log_2_(*p*). Both, for visibility of the text and to indicate an additional p-value cut-off the text (t-values) is plotted in white if *p ≤ p*_text_. The bottom legend shows 5 *p*-values (text: log_10_(*p*)) and the resulting field and text colors. Here *p*_min_ = 10*^−^*^10^ and *p*_text_ = 10*^−^*^5^.

**Figure S14.**
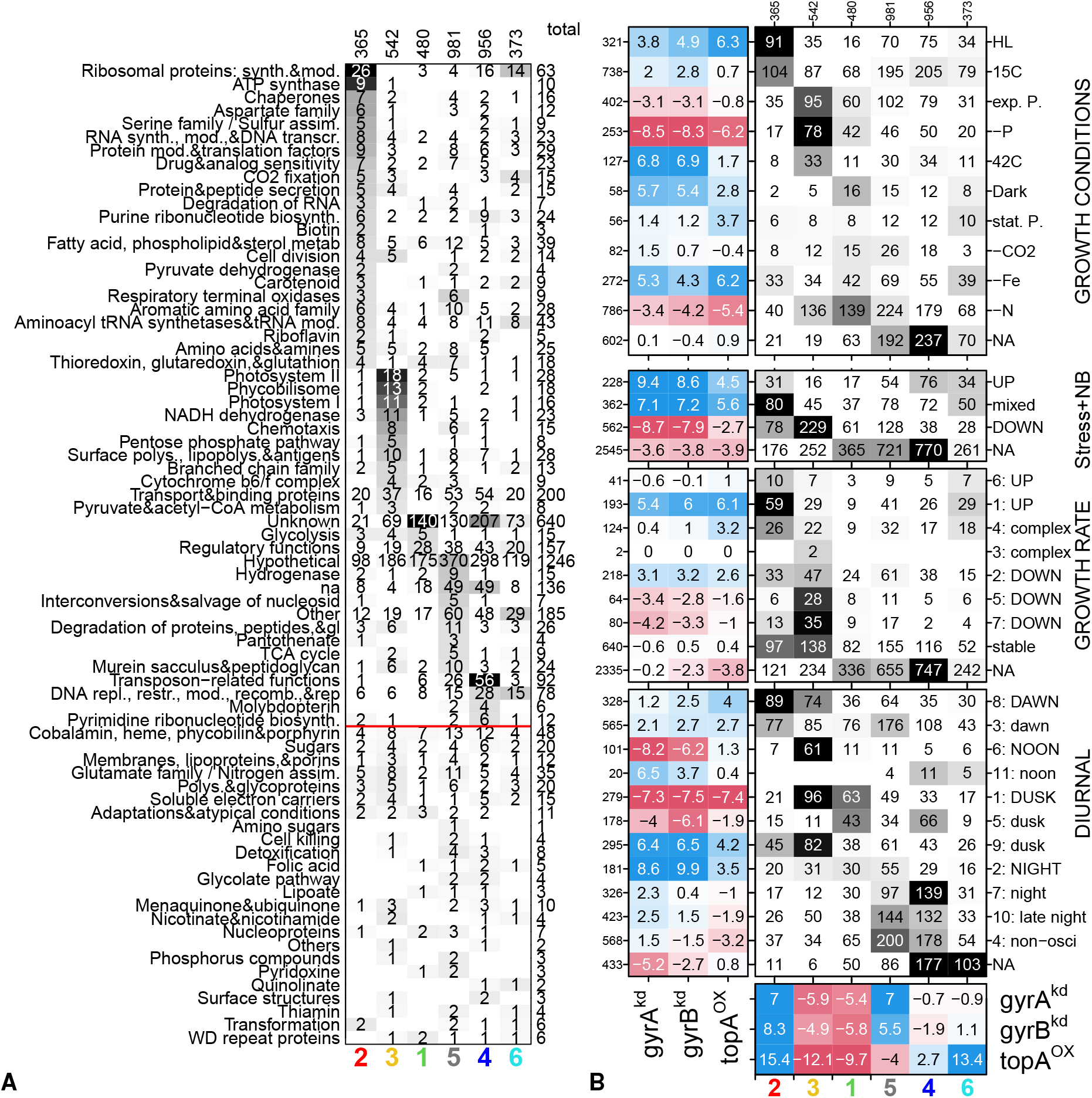
CyanoBase Category Analysis of Co-Expressed Cohorts. **A:** Sorted enrichment profile of functional category annotations as in Figure 5E (colored with *p*_min_ = 10*^−^*^10^ and *p*_text_ = 10*^−^*^5^) but sorted at *p*_sort_ = 0.1. All categories below the red line had only *p > p*_sort_ and are unsorted. Some abbreviations of the original annotation terms are used for readability of the plot: synth. – synthesis, mod. – modification, repl. – replication, transcr. – transcription, recomb. – recombination, restr. – restriction, s. – saccharides, assim. – assmilation, & – and. **B:** Overlap enrichement and t-test profiles with clusterings as Figure 6C but for additional gene classifications from other publications; from top to bottom: experimental `GROWTH CONDITIONS` with maximal expression of transcription units from Kopf *et al.* [50], stress and novobiocin (Stress + NB) treatment (same as in Fig. 6C) from ref. [4], the original non-collapsed clustering of protein abundance level response to `GROWTH RATE` from Zavrel *et al.*. [14], and a clustering of a `DIURNAL` transcriptome data set from the supplemental material of Lehmann *et al.* [34].

**Figure S15.**
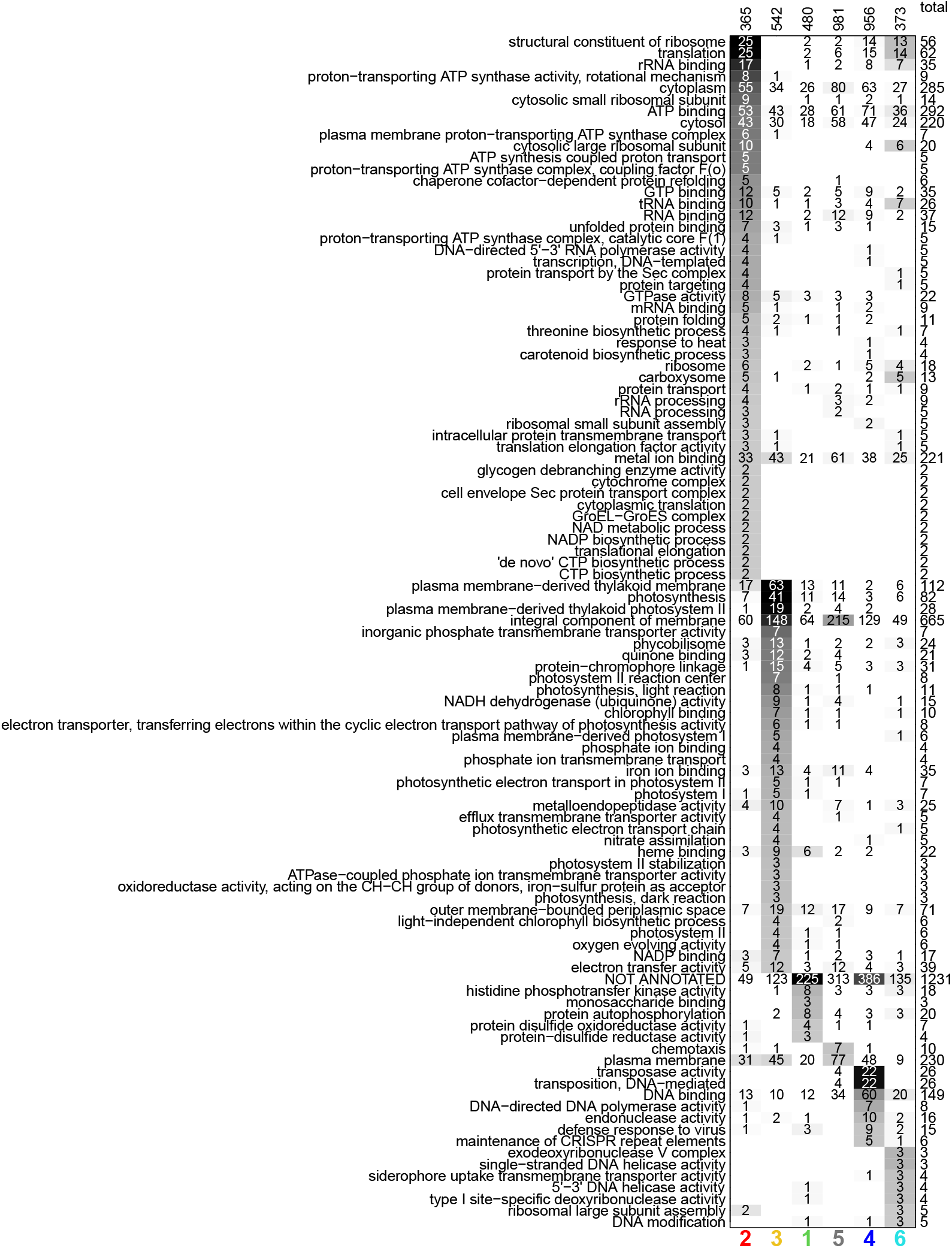
GO Analysis of Co-Expressed Cohorts. Sorted enrichment profile as in Figures 5E and S14 A (colored with *p*_min_ = 10*^−^*^10^ and *p*_text_ = 10*^−^*^5^) but for Gene Ontology (GO) terms, downloaded from the `UniProt` database (2021-03-20, organism:1111708). Rows are cut and sorted along columns at *p*_sort_ = 0.01.

**Figure S16.**
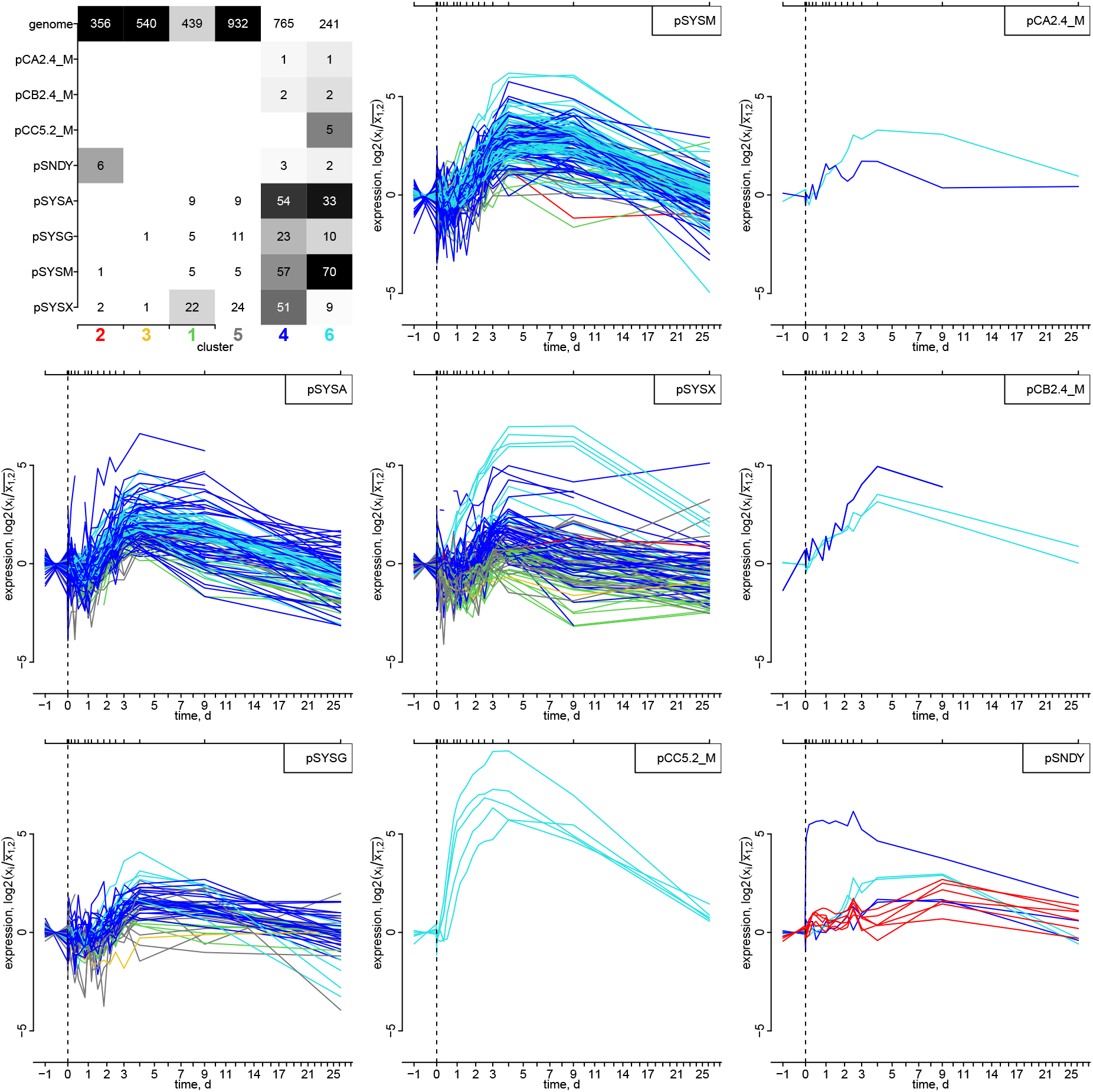
Transcriptome Time Series – Plasmids. Top left panel: Enrichment profile of time series clusters with the locations on the chromosome, one of the seven endogenous plasmids, or our construct pSNDY [36] (Table 1). All other panels show the temporal transcript abundance profiles for the coding genes of each plasmid (see top right legends for plasmid names); each transcript is colored according to its cluster label.

**Figure S17.**
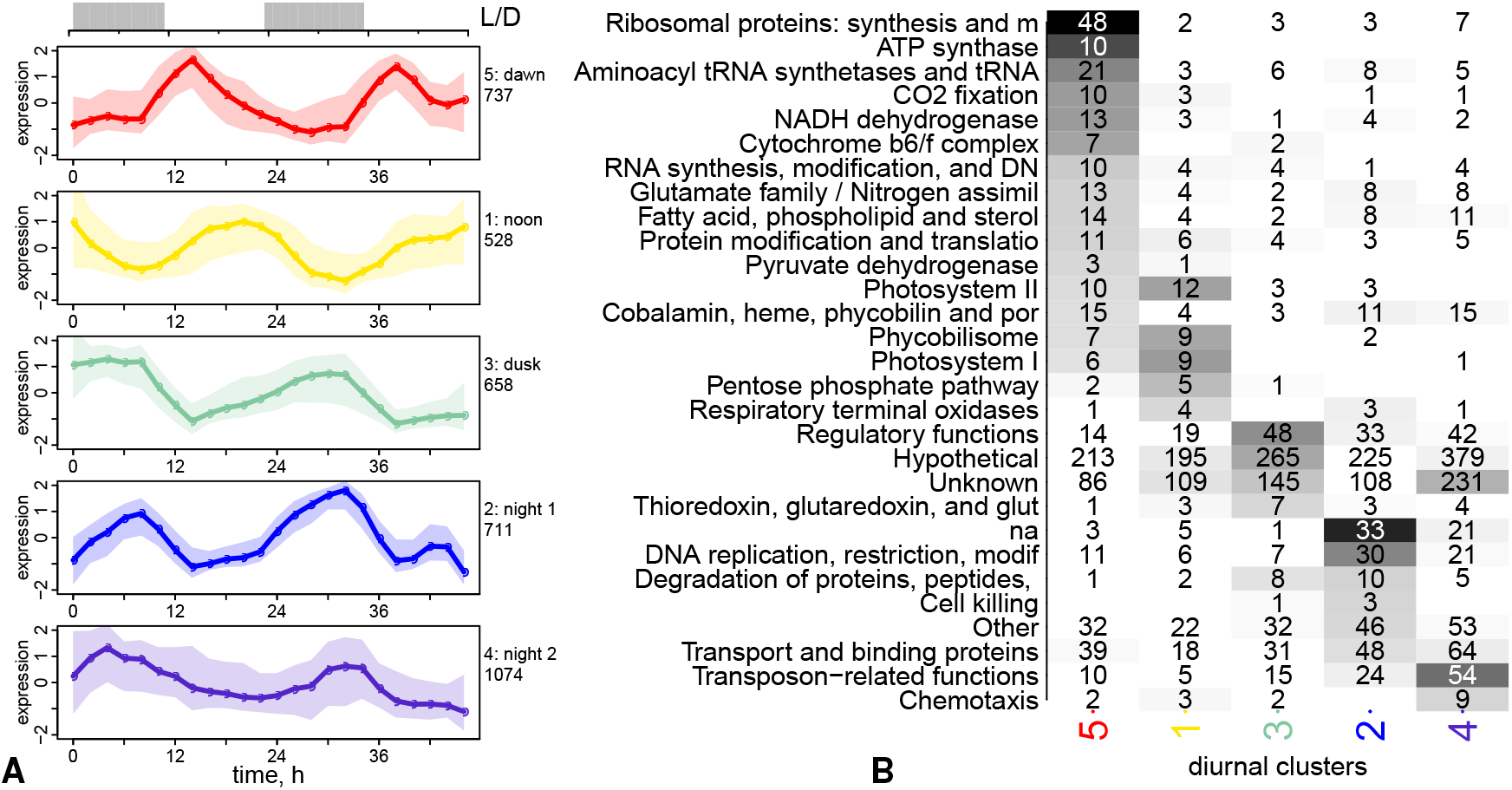
Diurnal Co-Expression Cohorts. Clustering of diurnal transcriptome data from ref. [48] into 5 co-expression cohorts, see Methods for details. **A:** Cluster medians of the normalized (to mean 0) expression values with an additional moving average over 3 samples. Transparent ranges show the 10% and 90% quantiles, *i.e.* they encompass 80% of all values in a cluster. Cluster labels and sizes (number of genes) are indicated on the right y-axis. The gray and white bars on the top indicate dark and light phases of the experiment. **B:** Enrichment profiles of co-expressed cohorts with CyanoBase functional categories as for Figure 5E (colored with *p*_min_ = 10*^−^*^10^ and *p*_text_ = 10*^−^*^5^), but cut and sorted at *p*_sort_ = 0.05.

**Figure S18.**
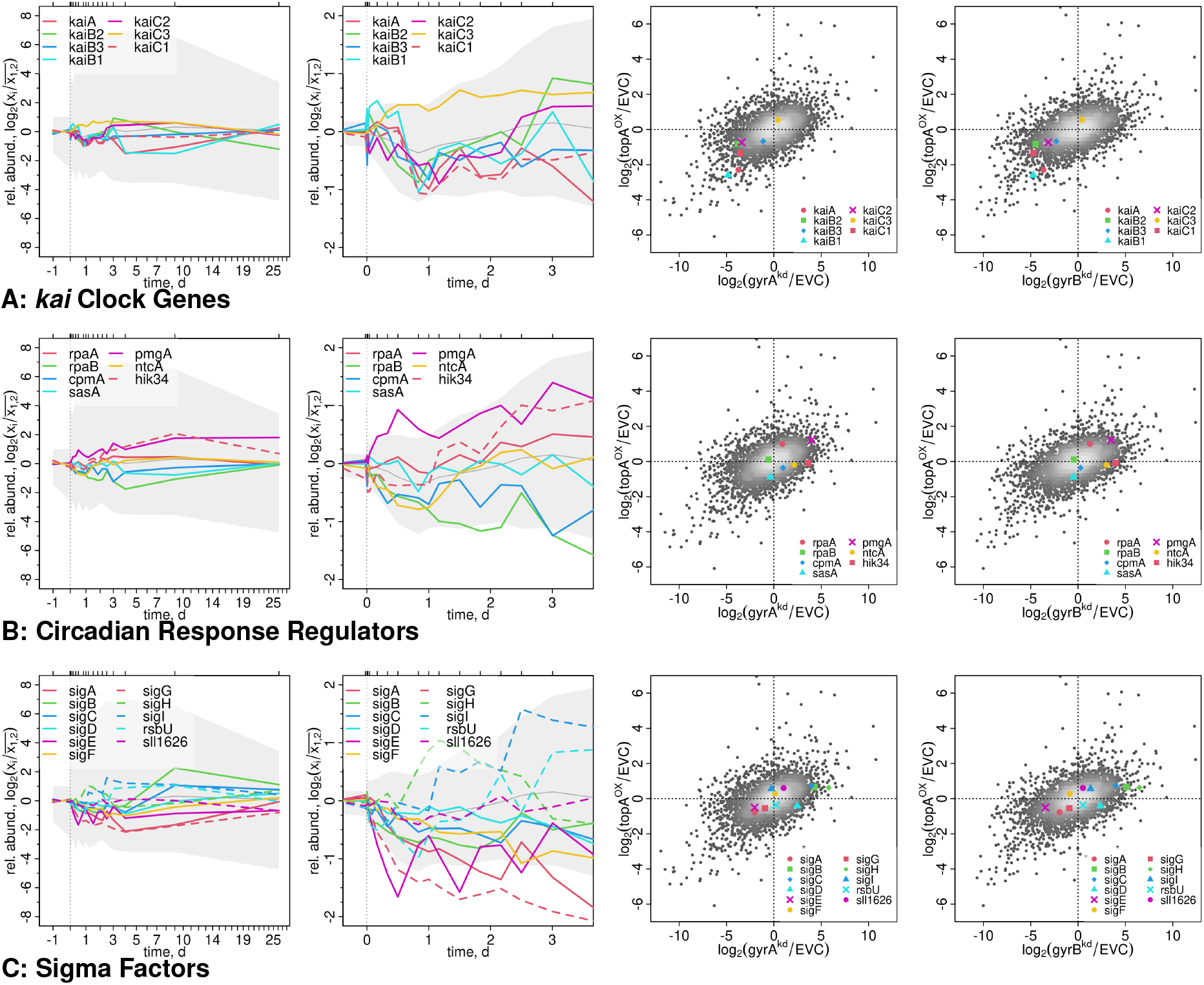
Genes of Interest: Regulators. Transcript abundance profiles of a selected groups of genes; from left to right: the full time-course, a zoom on the first three days, and expression changes with respect to the EVC in the endpoint experiments (as in Fig. 4D,E). The gray background in the time series plots shows the 92.5 % and 7.5 % quantiles of all data as a reference, the gray dots in the endpoint experiments are all other genes and the gray scale indicates local density. Some genes with strong response are specifically mentioned in the following. **A, Clock Genes:** During the first half day post-induction *kaiB1* was upregulated, then quickly downregulated with most other *kai* genes. Only *kaiC3* was upregulated, reflecting the pattern of the RB/dawn cohort, and then remained slightly over-expressed until the last sampled time point. At 3 d *kaiB2* and *kaiC2* were upregulated. **B, Signaling:** The circadian clock output regulator *rpaA* was only slightly upregulated, while its paralog *rpaB* was downregulated. **C, Sigma Factors:** Most sigma factors [55] were downregulated, except for the group 3/4 factors *sigH*, peaking at 1 d, followed by *sigI* (2 d–3 d), and a late peak (10 d, when cells were already enlarged) of the group 2 and stress-response sigma factor *sigB*.

**Figure S19.**
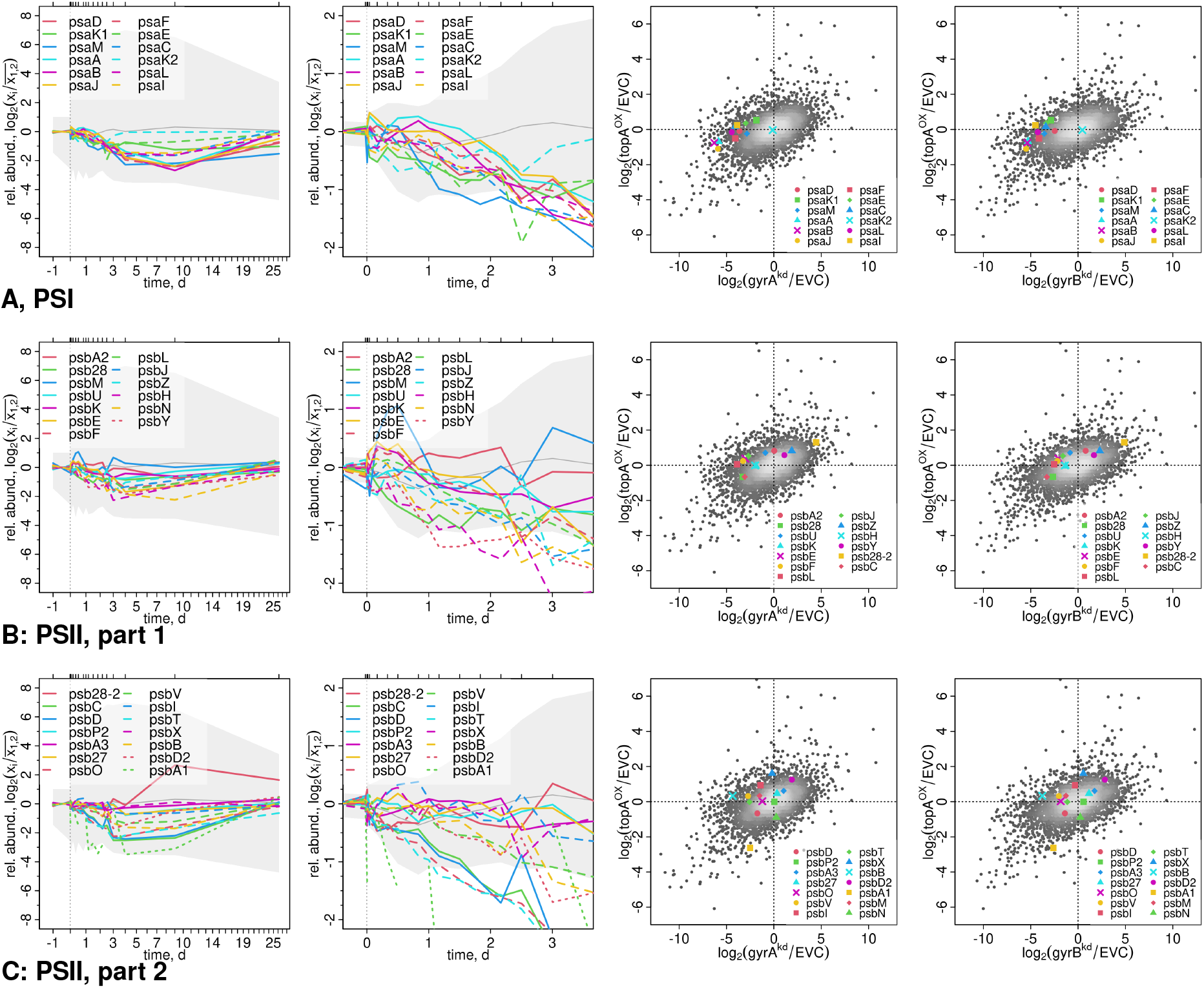
Genes of Interest: Photosystem I & II. As Figure S18 but for genes encoding for the photosystems. **B/C, PSII:** *psb28-2*: photosystem II reaction center protein Psb28 homologue, extrinsic protein of photosystem II, *psb28-1* but not *psb28-2* was required in PSII recovery after high-light [79].

**Figure S20.**
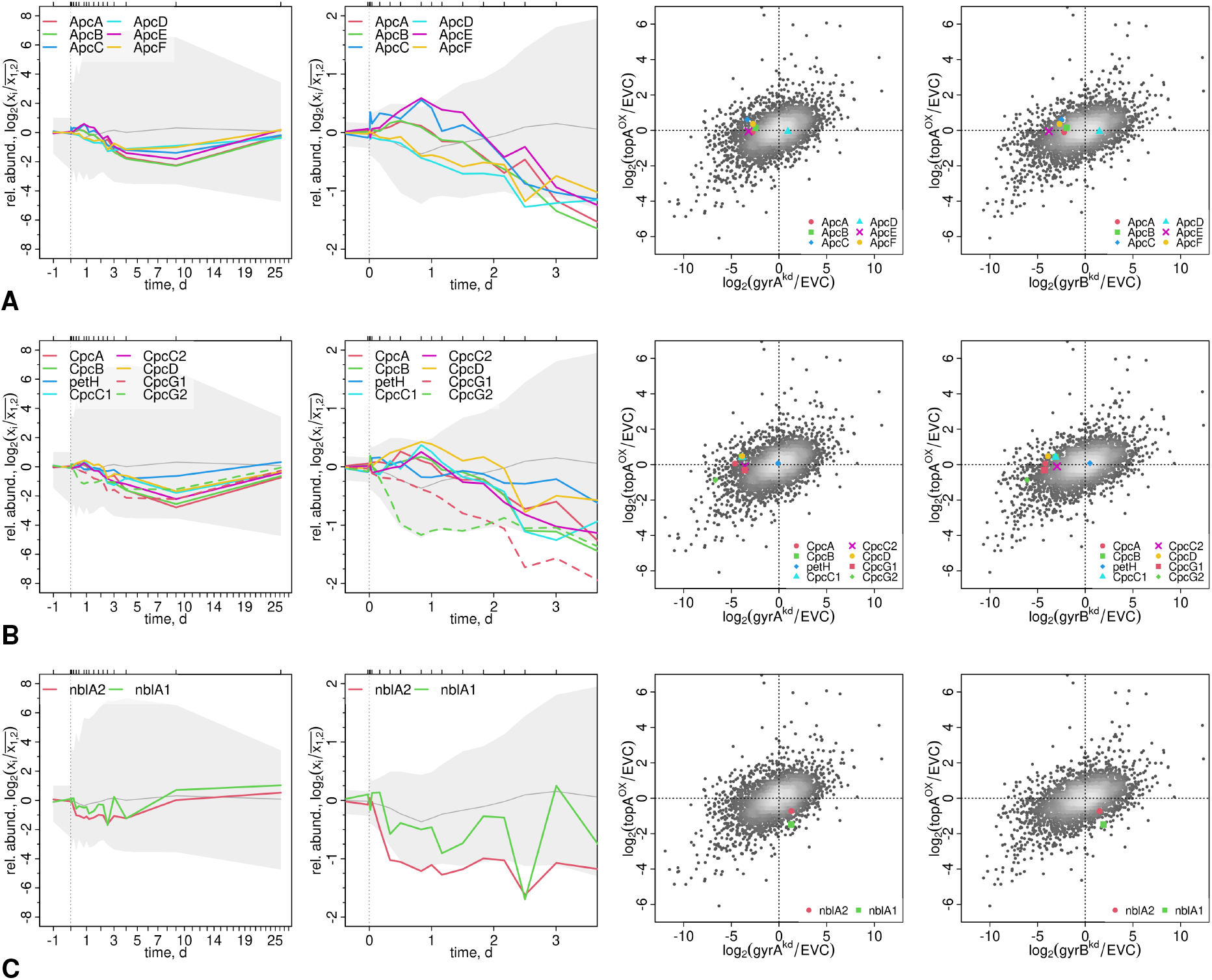
Genes of Interest: Phycobilisome. As Figure S18 but for genes encoding for the phycobilisome. Phycobilisome genes were downregulated in all experiments, except for the topA^OX^ endpoint measurements (right panels).

**Figure S21.**
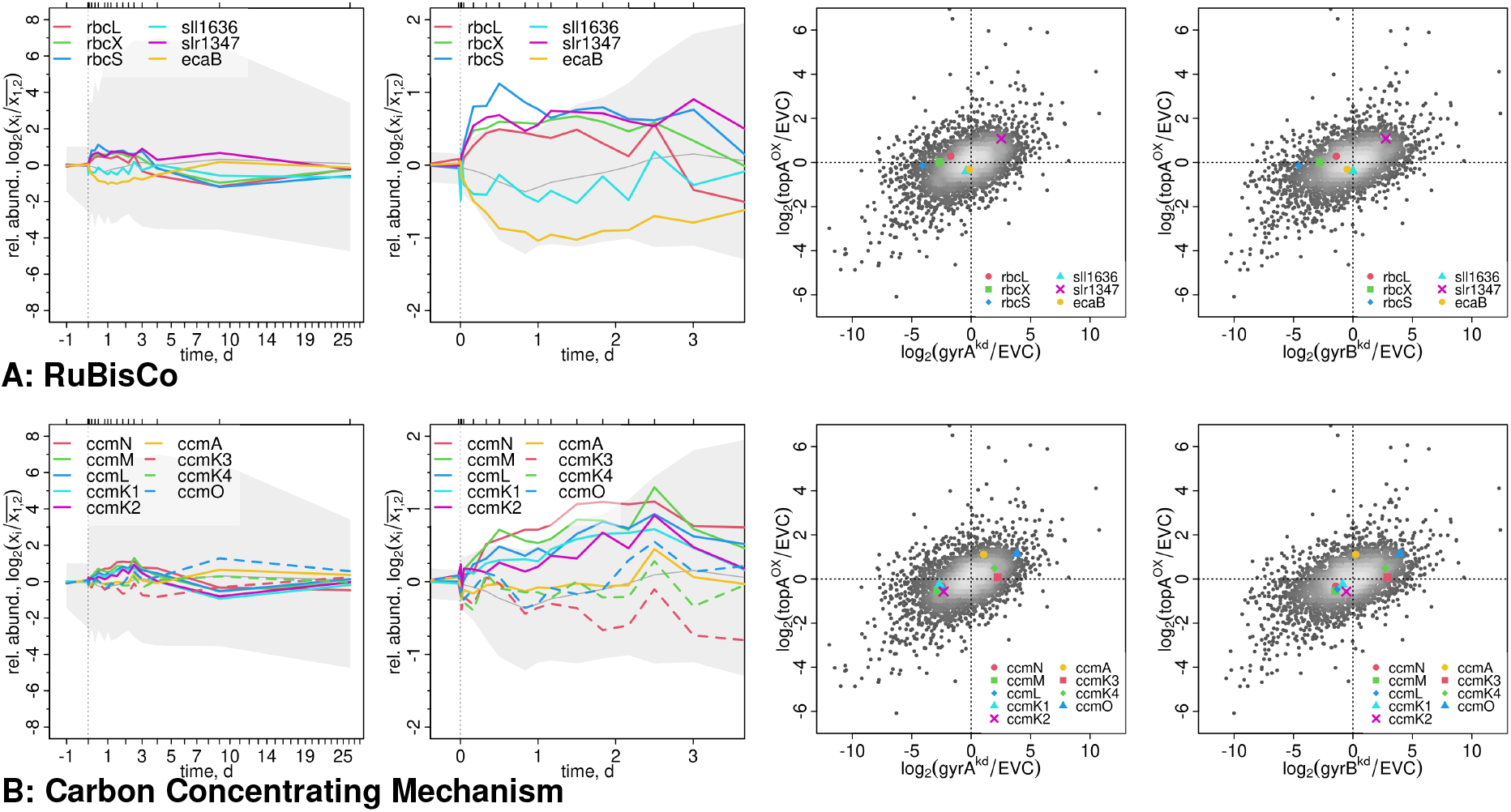
Genes of Interest: RuBisCo, Carbon Concentrating Mechanism. As Figure S18 but for genes encoding for RuBisCo and carbon concentrating mechanisms. **A, RuBisCo:** large and small subunits of RuBisCo (*rbcL/S* and the assembly factor (*rbcX* were all upregulated. *ecaB*: carbonic anhydrase; *slr1347*: carbonic anhydrase; *sll1636*: gamma carbonic anhydrase. One carbonic anhydrase *slr1347* was upregulated, while the other two were downregulated.

**Figure S22.**
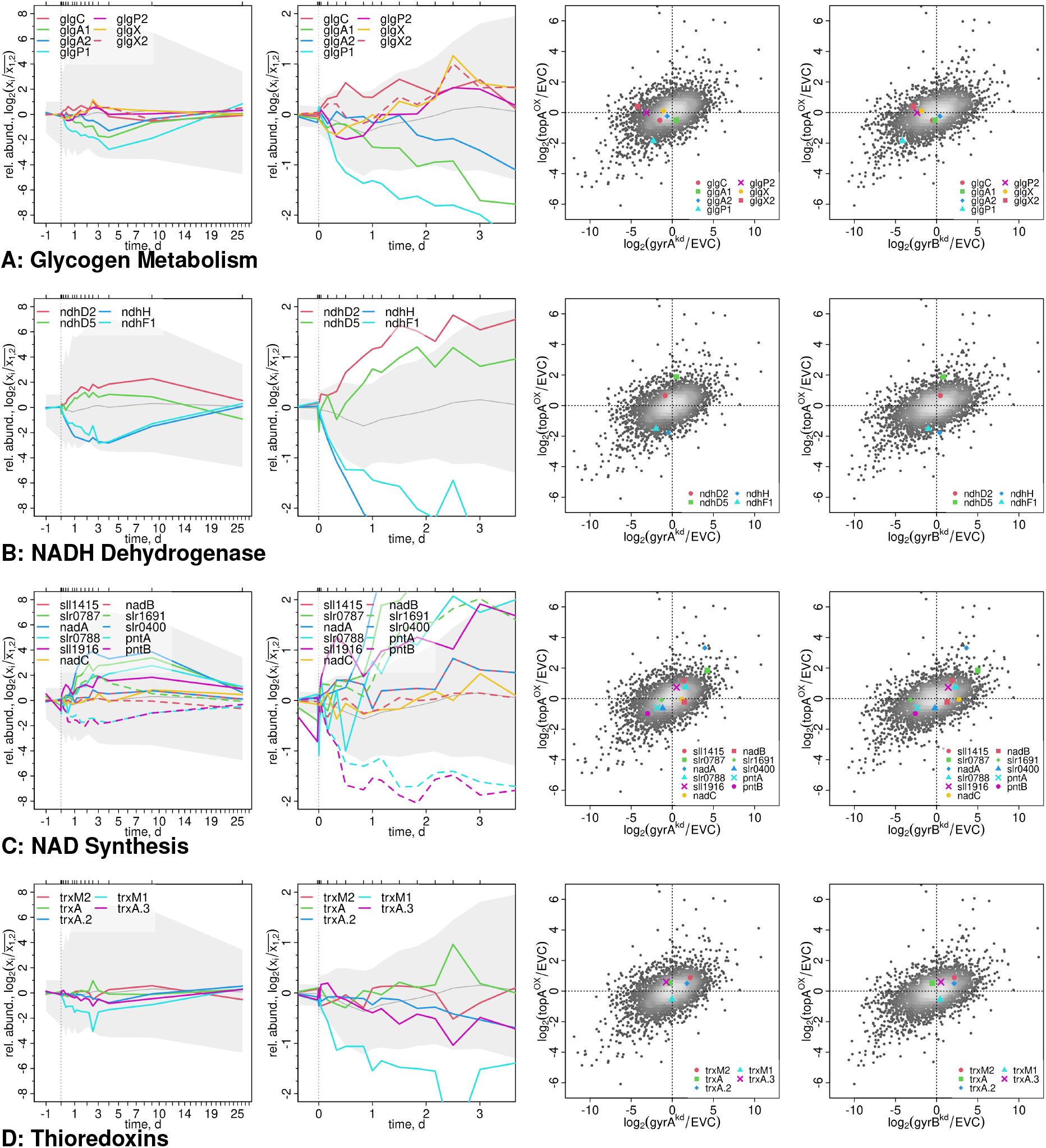
Genes of Interest: Metabolism. As Figure S18 but for genes that encode for metabolic enzymes/pathways. **A, Glycogen:** The glycogen degrading enzyme *glgP1* [80] was strongly downregulated, but *glgP2* upregulated at 3 d. The glucose-1-phosphate adenylyltransferase *glgC* was upregulated with the RB/dawn cohort, but both glycogen synthase genes, *glgA/A2*, where downregulated. The glycogen debranching enzymes, *glgX/X2*, were both upregulated at 3 d. **B, NADH dehydrogenase:** selected genes from the NADH dehydrogenase complex that showed a strong response, notably, only in the topA^OX^ strain but not the gyr^kd^ strains; their annotations are *ndhD2 (slr1291)*: electron transfer from NADH to plastoquinone; *ndhD5 (slr2007)*: Na-proton antiporter; *ndhH/ndhF1 (slr0261/slr0844)*: NAD(P)H-quinone oxidoreductase subunit H (chain 7) and chain 5. Notably, a knock-out of *pgmA* had impaired growth in photomixotrophic conditions, with frequent revertant mutations in NADH dehydrogenase subunits [56]. **C, NAD metabolism:** *nadA (sll0622)*: quinolinate synthasegenes involved in NAD metabolism, *pntA/B (slr1239/slr1434)*: transhydrogenase, transfers H from NADPH to NAD. **D, Thioredoxins:** The thioredoxin *trxM1* is strongly down-regulated, while *trxA* has a short peak at 2 d–3 d.

**Figure S23.**
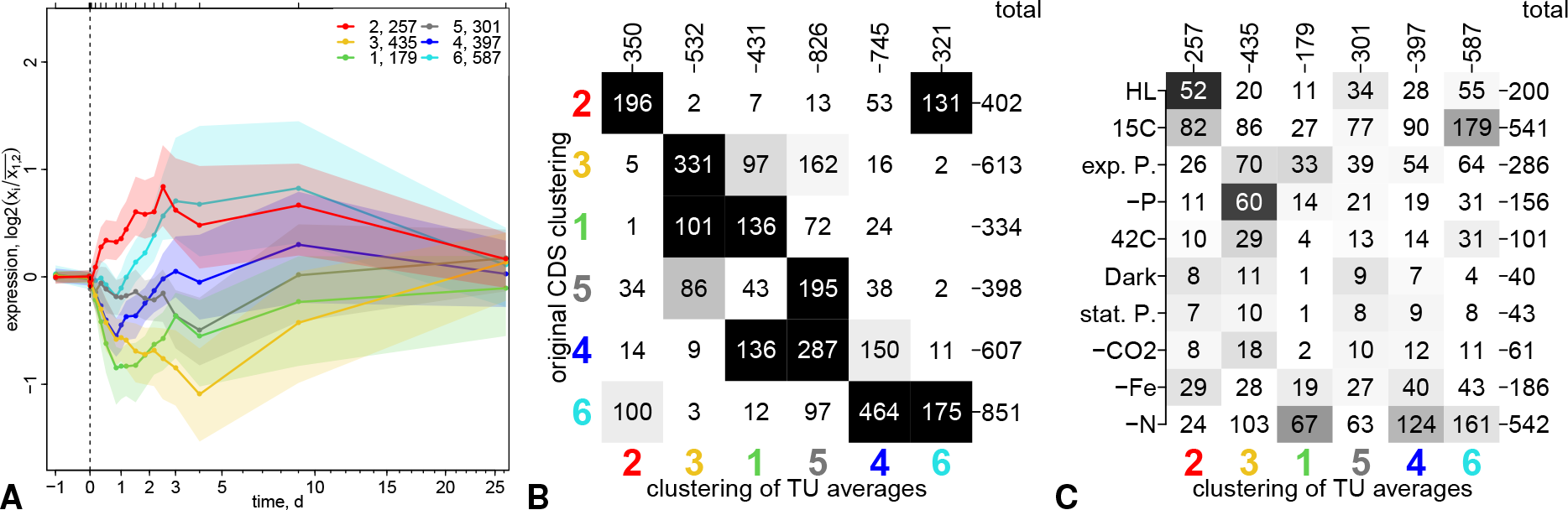
Transcription Start Site Analysis. A: Clustering of transcription units (TU) defined by [50]. Average expression was calculated for all TU from the expression of coding genes they encompass (*via* the “Sense.tags” column of the original data set), and the resulting TU time-series was clustered by k-means, using cluster centers from the CDS clustering (Fig. 5) and identical time-series processing. **B:** enrichment profile of the original CDS clustering (y-axis) with the TU-based re-clustering; colored with *p*_min_ = 10*^−^*^10^ and *p*_text_ = 10*^−^*^5^ and with the original order. **C:** enrichment profile of the time-series clusters with the original TU classification by condition of maximal expression (column “Max.cond.”) by [50]; colored with *p*_min_ = 10*^−^*^10^ and *p*_text_ = 10*^−^*^5^ but using the original orders. Note, that many CDS were re-assigned to different clusters but the overall pattern is reproduced.

## Footnotes

1 ^1^*** ref. [35] is a preprint by our group at bioRxiv: https://doi.org/10.1101/2021.07.26.453679 ***

2 ^2^*** ref. [35] is a preprint by our group at bioRxiv: https://doi.org/10.1101/2021.07.26.453679 ***

## Notes

### Competing Interest Statement

The authors have declared no competing interest.

### Summary of Updates

* Results and Discussion: References and more discussion added for hypernegative supercoiling of plasmids, downstream gyrase activity and binding sites (total RNA and rRNA analysis), and night-specific ppGpp regulation (bifurcation of response). The final paragraph on genomic vs. plasmid DNA replication was simplified. * The conclusion was shortened and clarified.

